# Super elongation complex as a targetable dependency in diffuse midline glioma

**DOI:** 10.1101/2020.01.22.913244

**Authors:** Nathan A. Dahl, Etienne Danis, Ilango Balakrishnan, Dong Wang, Angela Pierce, Faye M. Walker, Ahmed Gilani, Natalie J. Serkova, Krishna Madhavan, Susan Fosmire, Adam L. Green, Nicholas K. Foreman, Sujatha Venkataraman, Rajeev Vibhakar

## Abstract

Mutations in the histone 3 gene (H3K27M) are the eponymous drivers in diffuse intrinsic pontine gliomas (DIPGs) and other diffuse midline gliomas (DMGs), aggressive pediatric brain cancers for which no curative therapy currently exists. The salient molecular consequence of these recurrent oncohistones is a global loss of repressive H3K27me3 residues and broad epigenetic dysregulation. In order to identify specific, therapeutically targetable epigenetic dependencies within this disease context, we performed an shRNA screen targeting 408 genes classified as epigenetic/chromatin-associated molecules in patient-derived DMG cultures. This approach identified AFF4, the scaffold protein of the super elongation complex (SEC), as a previously-undescribed dependency in DMG. Interrogation of SEC function demonstrated a key role for maintaining DMG cell viability and clonogenic potential while promoting self-renewal of DMG tumor stem cells. Small-molecule inhibition of the SEC with the highly-specific, clinically relevant CDK9 inhibitors atuveciclib and AZD4573 restores regulatory RNA polymerase II pausing, promotes cellular differentiation, and leads to potent anti-tumor effect both *in vitro* and in patient-derived xenograft models. These studies present a biologic rationale for translational exploration of CDK9 inhibition as a promising therapeutic approach in a disease which currently has no effective medical therapies.

## Introduction

Diffuse intrinsic pontine glioma (DIPG) is a uniformly fatal pediatric brain tumor with extraordinarily limited treatment options. The median survival for DIPG remains 11 months, with fewer than 5% of patients alive at 2 years from time of diagnosis(*1*). Curative surgery is not possible, radiation therapy provides only temporary relief, and chemotherapies have thus far proven wholly ineffective(*2, 3*). The current World Health Organization (WHO) diagnostic classification of diffuse midline glioma (DMG) is now categorically defined by the presence of the H3K27M mutation(*4*), and the clinical relevance ascribed to this genetic lesion highlights its unifying role as the biologic driver across a range of midline glial tumors involving the thalamus, cerebellum, brainstem (DIPG), and spinal cord(*5–7*). The presence of the H3K27M mutation in these tumors conveys a uniformly dismal prognosis regardless of tumor location or histologic grade(*6, 8*). There exists a compelling need to understand the mechanisms by which H3K27M-mediated epigenetic perturbation disrupts normal neuroglial development in order to develop targeted therapies to benefit this high-risk subset of patients.

Mutations to the *H3.3*, *H3F3A*, or *HIST1H3B* histone genes have been identified in approximately 80% of pediatric DIPG(*9, 10*). This characteristic lysine-to-methionine (K27M) substitution leads to a profound global hypomethylation at H3K27-trimethyl-bound promotors and a subsequent loss of transcriptional repression at these loci(*11*). This observation has prompted prior preclinical investigations into the efficacy of transcriptional disruption in DMG, an effective strategy in many experimental models of transcriptionally addicted malignancies(*12–16*). Both chromatin targeting via bromodomain and extra-terminal (BET) family protein BRD4 inhibition using JQ1(*17–19*) and CDK7-mediated transcription initiation using THZ1(*20*) have demonstrated promise as druggable targets in patient-derived DMG cell cultures and xenograft models. These compounds, however, have proved challenging to adapt for clinical use. The relatively nonspecific approach of inhibiting genome-wide transcription may also impart off-target consequences given the comparatively restricted transcriptomic changes observed in H3K27M-mutant tumors(*21*). In this study, we set out to identify specific epigenetic regulators which represent critical cellular dependencies within the context of an H3K27M+ transformed cell state, to understand how they might mediate the altered developmental phenotype observed in DMG cell populations, and to determine whether these represented vulnerabilities that might be amenable to disruption using clinically relevant compounds.

## Results

### Epigenome screening identifies AFF4 as an epigenetic dependency in DMG

In order to identify specific epigenetic machinery that acquires an aberrant or critical function in the presence of the H3K27M mutation, we performed a functional shRNA screen using 4188 unique shRNA sequences targeting 408 genes classified as epigenetic or chromatin-associated molecules (4-12 individual shRNA sequences per gene). SU-DIPG4 cells were stably transduced in triplicate via a pooled lentiviral vector system and passaged for 21 days before isolating genomic DNA from surviving cells. Incorporated shRNA sequences were quantified in comparison to an immediate post-transduction sampling, with depleted sequences representing candidate essential genes (**Fig 1A**). shRNAs were ranked by the observed decrease in cell viability (**Fig 1B**), identifying 47 candidate genes with a fold change <0.8 and p <0.05 (**Fig 1C** and **Table S1**). These hits were enriched in genes involved in chromatin reading and writing functions with previously reported biologic relevance in DMG, including histone deacetylases, histone methyltransferases, polycomb repressive complex 1 members, and bromodomain family proteins. We prioritized genes which had not been previously characterized within a DMG model and which had related pharmacologic inhibitors in clinical use or preclinical development. Secondary validation of hits using individual shRNA constructs was performed in two additional H3K27M+ DMG cell lines (SF8628 and SF7761)(**Fig 1D**). Analysis of this screening approach identified depletion of AFF4 as a novel lethality in DMG.

**Figure 1.**
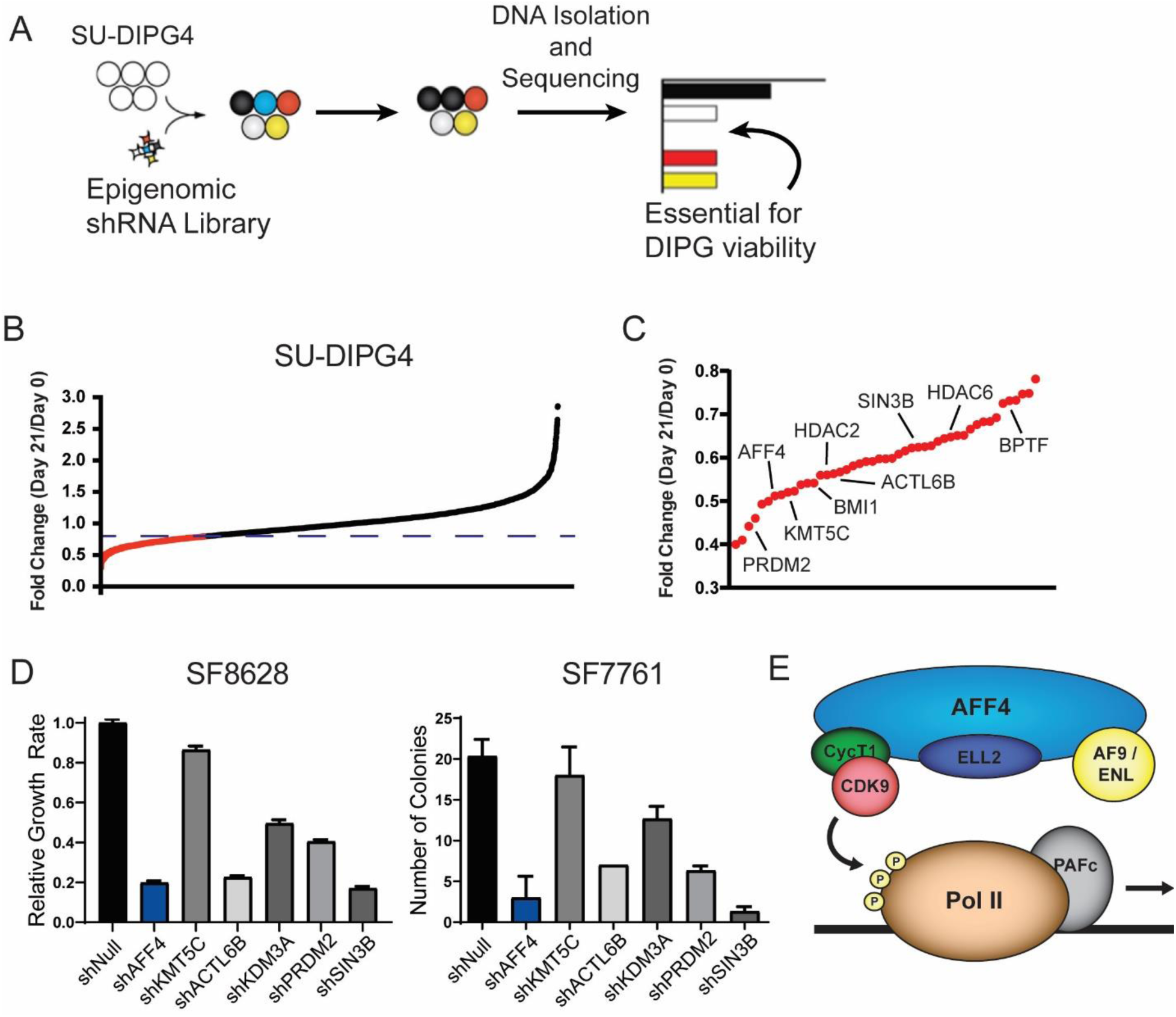
Functional epigenome-wide screen identifies AFF4 as an epigenetic dependency in H3K27M+ DMG. **A.** Pooled epigenomic shRNA library is transduced into SU-DIPG4 cells, and genomic DNA is isolated and sequenced to identify individual shRNA amplification or loss. **B.** Aggregate normalized sequencing data showing depleted shRNAs (fold change </= 0.8 in red). **C.** Leading edge of depleted shRNAs ordered by fold change. **D.** Secondary validation of screening hits by growth rate (SF8628, left) or colony formation (SF7761, right). Data displayed as mean ± SD. **E.** Model of the SEC in which CDK9 phosphorylates the carboxyl-terminal domain of RNA Pol II, resulting in release from promotor-proximal pause site.

The AFF4 protein forms the scaffold to which other SEC subunits bind, including AF9/ENL, ELL2, and the CycT1-CDK9 pairing collectively termed positive transcription elongation factor b (P-TEFb) (*22, 23*). The SEC acts as a positive regulator of the release of RNA polymerase II (Pol II) from promoter-proximal pausing, a key regulatory step conserved across metazoan development (*24, 25*). Phosphorylation of Pol II C-terminal domain (CTD) at the serine 2 position by P-TEFb, the catalytic subunit of the SEC, allows for release of Pol II from this paused state and for productive transcriptional elongation to occur (*26, 27*) (**Fig 1E**). SEC mediation of Pol II pausing release is vital for the dynamic induction of developmentally-regulated genes in response to cellular differentiation signals and can be misregulated in states of cellular transformation (*23, 28–30*). The role of SEC misregulation in DMG is not known.

### SEC is critical for DMG cell proliferation and mediates stem-like phenotype

In order to interrogate the role of the SEC in DMG, we employed two nonoverlapping shRNA lentiviral constructs to deplete AFF4 (shAFF4) in three H3K27M-mutant DIPG (SU-DIPG4, HSJD-DIPG007, SF8628) and one H3K27M-mutant thalamic DMG (BT245) cell lines (**Fig 2A** and **Fig S1A**). We first examined DMG cell clonogenic potential by performing colony focus assays in SU-DIPG4 and HSJD-DIPG007 cells and observed a marked decrease in colony formation following shRNA-mediated reduction of AFF4 expression in comparison to non-targeting shRNA control (shNull) (**Fig 2B** and **Fig S1C-D**). We likewise found that AFF4 knockdown led to a significant decrease in proliferation across multiple patient-derived H3K27M+ cell cultures (**Fig 2C-D**). Though AFF4 is not known to have SEC-independent functions (*22, 23*), we confirmed its residence within an intact P-TEFb-containing complex in DMG by anti-AFF4 co-immunoprecipitation (**Fig 2E**). To further support that our observations reflected a complex-dependent phenotype, we used a similar lentiviral shRNA approach to deplete CDK9, the catalytic subunit of the SEC, in HSJD-DIPG007, SF8628, and BT245 cells (**Fig S1B**). Following transduction with both nonoverlapping shCDK9 constructs, comparable decreases in cell proliferation were observed (**Fig 2F**).

**Figure 2.**
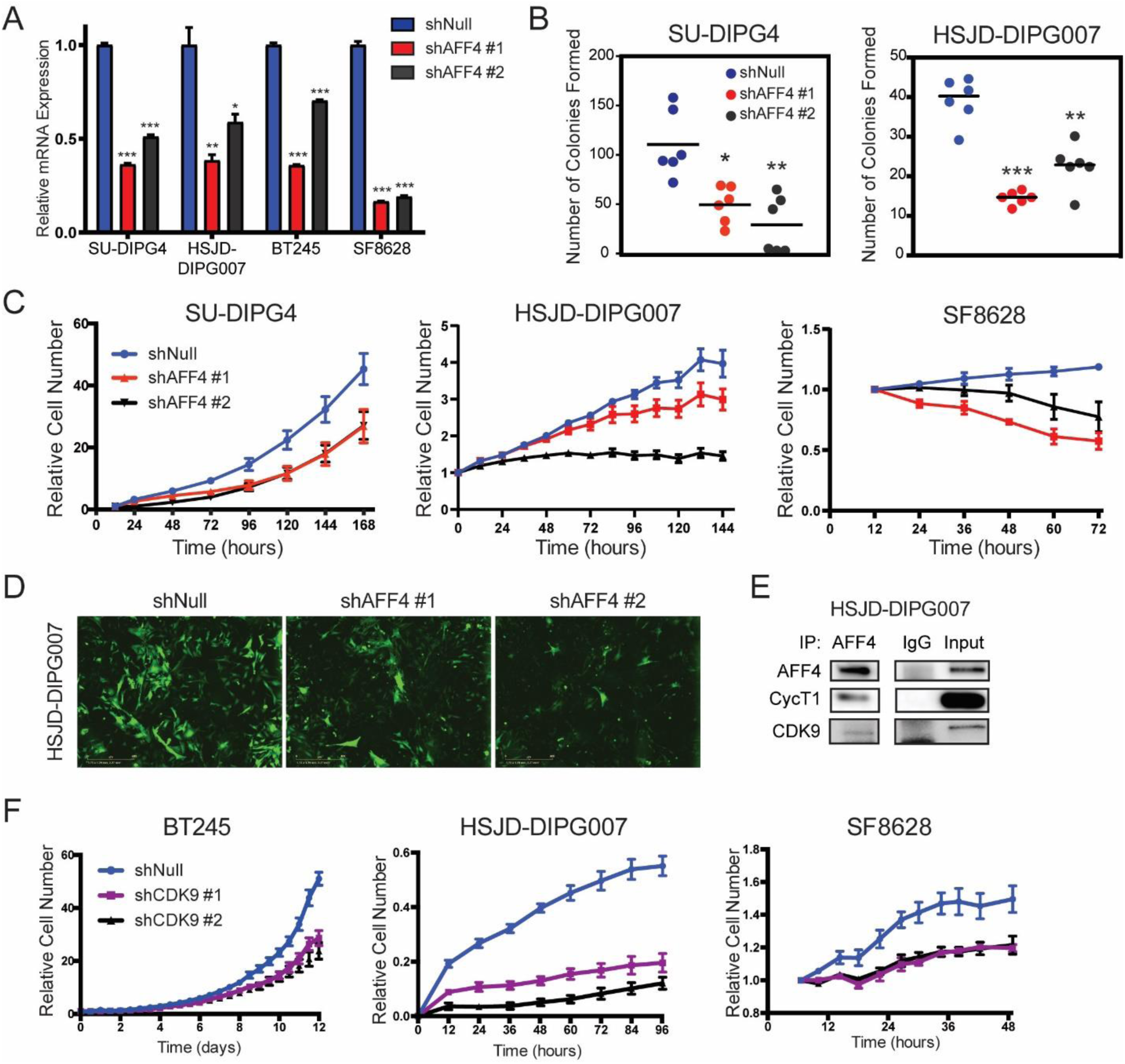
SEC is critical for DMG cell proliferation and clonogenic potential. **A.** Decrease in AFF4 mRNA levels in SU-DIPG4, HSJD-DIPG007, BT245, and SF8628 cell lines following transduction with two non-overlapping shRNA lentiviral constructs in comparison to shNull control (two-tailed t-test, *p<0.05, **p<0.001, ***p<0.0001). **B.** Clonogenic potential following shAFF4 knockdown as measured by adherent colony formation (two-tailed t-test, *p<0.05, **p<0.001, ***p<0.0001). **C.** Rate of adherent growth in SU-DIPG4, HSJD-DIPG007, and SF8628 cells treated with shNull or shAFF4. Data are displayed as mean ± SEM. **D.** Representative fluorescent live-cell imaging from HSJD-DIPG007 cells following respective lentiviral transduction. **E.** Co-immunoprecipitation with either anti-AFF4 or IgG control antibodies followed by immunoblot for P-TEFb member proteins. **F.** Rate of growth in BT245, HSJD-DIPG007, and SF8628 cells treated with shNull or shCDK9 constructs. Data are displayed as mean ± SEM.

To identify the transcriptional programs under SEC regulatory control in H3K27M-mutant DMG, we evaluated the effect of AFF4 depletion on the transcriptional landscape via RNA sequencing (RNA-seq). We compared BT245 cells transduced with either shNull or shAFF4 and identified 600 upregulated and 1391 downregulated genes (fold change </> 2 and p-value <0.01) (**Fig 3A**). Similar to previous studies (*31, 32*), we observed that SEC regulates *MYC* and Myc-dependent programs, with decreases in *MYC* and Myc targets such as *NPM2*, *CCND2*, and *FASN* seen following AFF4 knockdown. Upregulated Myc signaling may harbor relevance in a subset of DMGs (*33*), but the changes we observed downstream of Myc following AFF4 depletion were comparatively modest. In contrast, in addition to the expected defects in processive transcription, Gene Ontology (GO) and Gene Set Enrichment Analysis (GSEA) of differentially-expressed genes also demonstrated a higher-ranked enrichment in transcriptional programs involved in neuroglial differentiation (**Fig 3B-C**). We observed a decrease in expression of genes important for regulating progenitor cell state and self-renewal, including the Shh-responsive elements *PTCH2*, *GLI1*, and *SIX5*. Reciprocal increases in markers of neuroglial differentiation, such as *GFAP*, *NGFR*, and *NRN1*, as well as mediators of proliferation and differentiation, including *CDKN1A* and *CDKN2B*, were likewise noted following depletion of AFF4 (**Fig 3D**). In order to validate these findings, we seeded both control and AFF4-depleted cells in neurosphere conditions, a surrogate environment for assaying self-renewal, and observed a significant decrease in both the frequency of neurosphere initiating cells (BT245, **Fig 3E**) and neurosphere growth (SU-DIPG4, HSJD-DIPG007, BT245, **Fig 3F-G**). Studies in *Drosophila* have described Pol II pausing release as a critical regulatory mechanism for developmental control genes, including tissue specification and CNS midline patterning (*34–36*). Multiple groups have shown that H3K27M-mutant DMG tumors are enriched in proliferative, stem-like cells with a relative paucity of mature, differentiated astrocytes or oligodendrocytes (*37*). Our data suggests that the SEC may play a role in maintaining this observed differentiation arrest.

**Figure 3.**
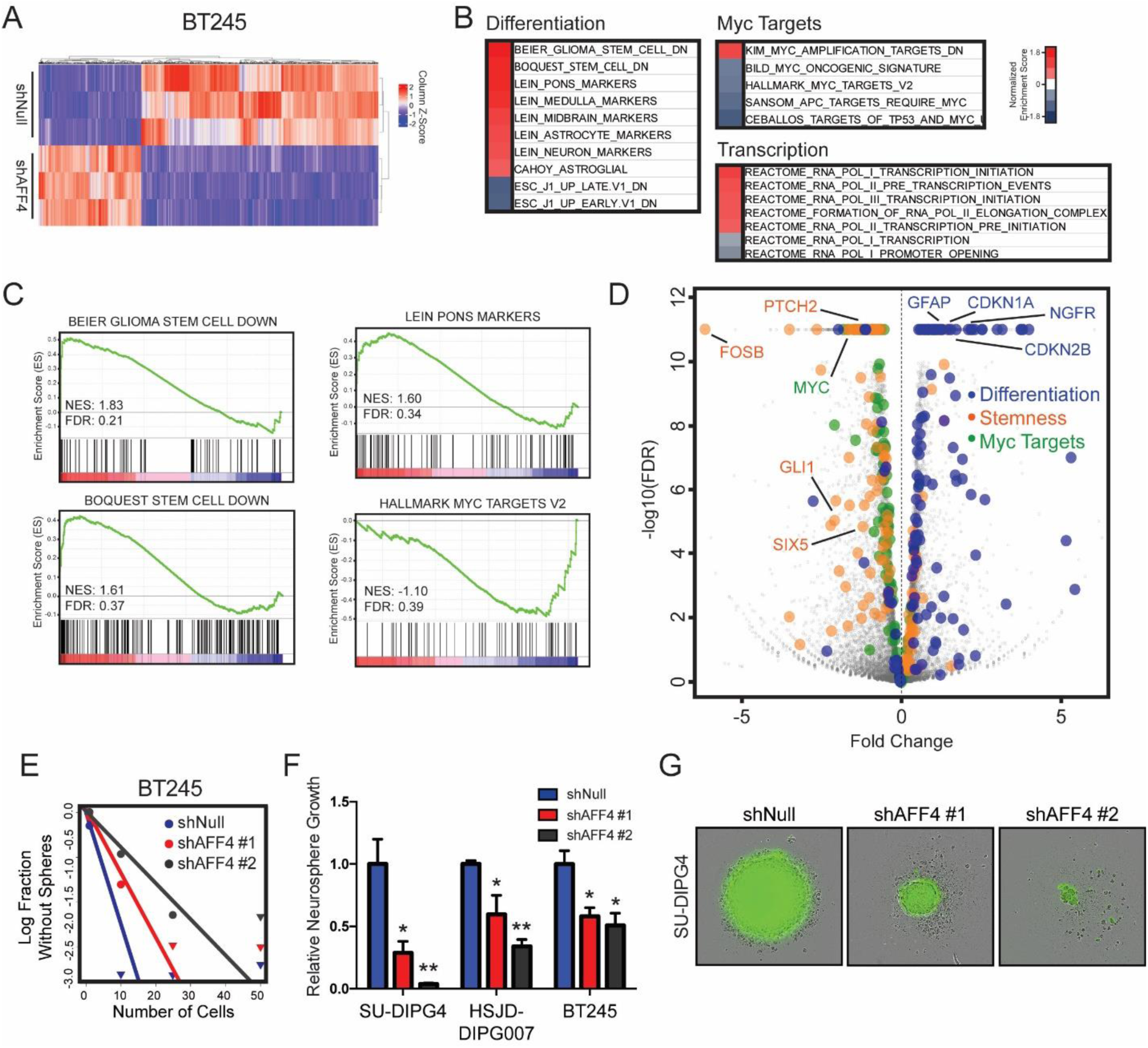
AFF4 depletion induces transcriptional profiles of cellular differentiation and decreases potential for self-renewal. **A.** Unsupervised hierarchical clustering of 1,671 differentially expressed genes from BT245 cells treated with either shNull or shAFF4 (n=3, fold change </> 2 and p-value <0.01). Gene set enrichment analysis results (**B**) and select enrichment plots (**C**) of differentiation- and Myc-associated gene sets, ordered by normalized enrichment score (NES). **D.** Differentially-expressed genes in differentiation-, stemness-, or Myc target-associated gene sets as defined by GSEA. Select illustrative genes with fold change </> 1 and p-value <0.01 are labeled. **E.** Neurosphere formation efficacy by extreme limiting dilution assay (χ^2^ shNull vs shAFF4 #1 p=0.035, vs shAFF4 #2 p=0.008). **F.** Normalized growth in neurosphere size of cells grown in suspension following transduction with shRNA constructs. Data are displayed as mean ± SEM. (two-tailed t-test, *p<0.05, **p<0.001). **G.** Representative images of neurospheres from SU-DIPG4 cells transduced with shRNA constructs containing a GFP reporter.

### CDK9 pharmacologic inhibition recapitulates phenotypic effects of SEC genetic depletion

AFF4 functions as a scaffold protein to which other SEC member units bind. It contains a long, intrinsically disordered N-terminal region responsible for interaction with other SEC subunits as well as a highly conserved C-terminal homology domain that interfaces with DNA and RNA (*38*). It is not, as of yet, directly targetable with clinically relevant compounds (*31*). CDK9, however, serves as the catalytic subunit of the SEC (**Fig 4A**) (*22, 23*). Dysregulated CDK9-mediated signaling has been described in numerous malignancies, prompting continued development of several CDK9 pharmacologic inhibitors (CDK9i) with increasing specificity (*23, 30, 39-41*). We sought to determine whether our findings from SEC genetic depletion could be replicated using a small-molecule therapeutic agent. We screened five commercially-available CDK9i compounds against a panel of six H3K27M-mutant cell lines and observed broad sensitivity in the sub- to low-micromolar range (**Fig 4B**). From this we chose two highly-selective agents currently in phase I clinical trials, atuveciclib and AZD4573 (median IC_50_ 769 nM and 9.5 nM, respectively), for further study (**Fig 4C**). We then performed RNA-sequencing following atuveciclib treatment in three H3K27M+ cell lines (SU-DIPG4, HSJD-DIPG007, and BT245) and identified approximately 1000 genes differentially expressed following CDK9 inhibition (range 918 – 1,447 genes, log fold change >1 and p < 0.05) (**Fig S2**). Gene Ontology meta-analysis revealed significant enrichment in gene sets involved in neuroglial differentiation programs, most notably synaptic signaling and cell projection or axonogenesis, along with signatures associated with cellular proliferation that were conserved across all three cell lines (**Fig 4D**). When compared within a given cell line (BT245), CDK9i treatment effectively phenocopied the transcriptomic effects of AFF4 shRNA knockdown on differentiation programs, with Metascape gene ontology analysis of common genes up- or downregulated by both genetic depletion and drug treatment again underscoring core signatures involved in regulating neuroglial morphogenesis (**Fig 4E** and **Table S2**). Atuveciclib treatment likewise diminished capacity for self-renewal in DMG cultures as assessed by both aldehyde dehydrogenase activity, a marker of the brain tumor initiating cell fraction with a given cell population (*42*) (**Fig 4F**), and neurosphere formation efficacy (**Fig 4G**).

**Figure 4.**
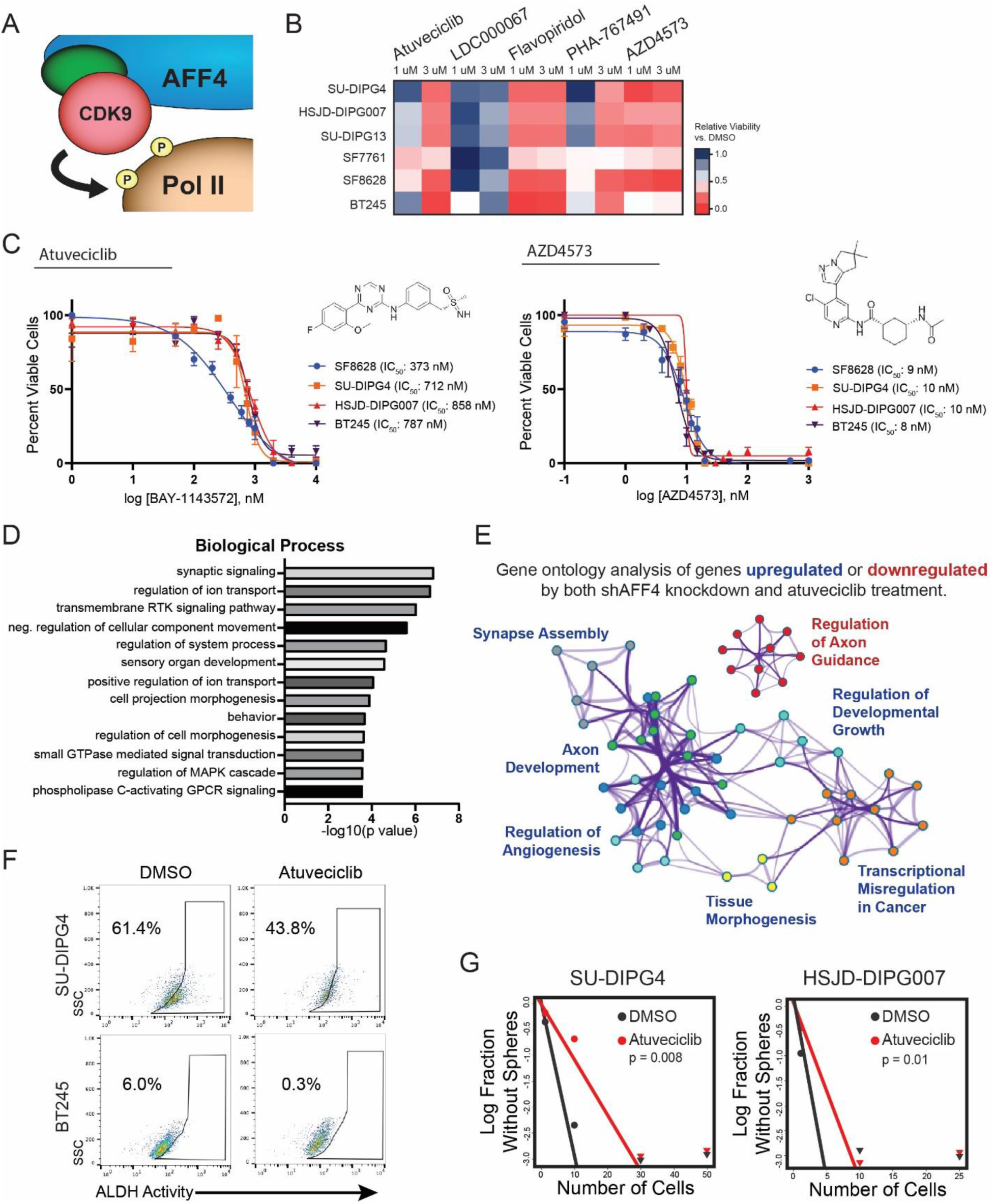
CDK9i treatment recapitulates phenotypic effects of SEC genetic depletion. **A.** CDK9 catalyzes phosphorylation of serine 2 in the carboxyl-terminal domain of RNA Pol II. **B.** Screen of five commercially available CDK9 inhibitors (atuveciclib, LDC000067, flavopiridol, PHA-767491, and AZD4573) against six H3K27M-mutant cell lines. Heatmap shown as relative cell viability vs DMSO. **C.** Sensitivity of DMG cultures to atuveciclib (left) and AZD4573 (right) as measured by half maximal inhibitory concentrations. **D.** Gene ontology terms significantly enriched across all three cell lines following atuveciclib treatment, ordered by -log10(p value). **E.** Metascape gene ontology network cluster constructed from common genes up- or downregulated by both shAFF4 knockdown and atuveciclib treatment. Each node denotes an enriched term, and networked clusters are indicated by different colors. **F.** Identification of brain tumor initiating cell fraction in SU-DIPG4 and BT245 cells by ALDH expression (ALDEFLUOR assay) demonstrates decrease in ALDH^+^ fraction following atuveciclib treatment compared to DMSO. **G.** Neurosphere formation efficacy by extreme limiting dilution assay following atuveciclib treatment of SU-DIPG4 (χ^2^ p=0.008) and HSJD-DIPG007 (χ^2^ p=0.01) cells.

**Figure 5.**
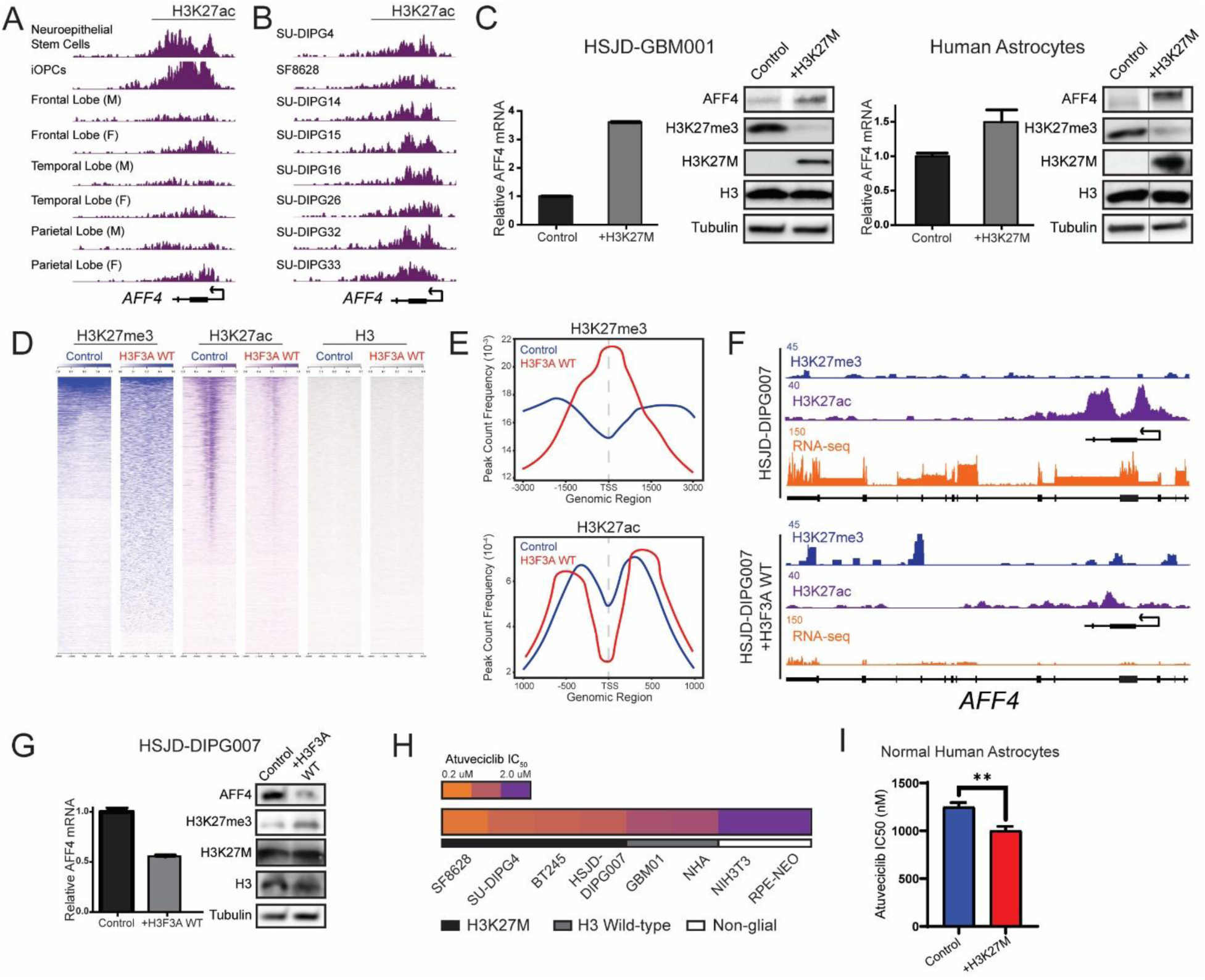
The H3K27M mutation alters epigenetic regulation of *AFF4* promoter. **A.** H3K27ac deposition at the *AFF4* promoter in neuroepithelial stem cells, OPCs, and differentiated brain regions (identical reads and scale for each sample tract). **B.** H3K27ac deposition at the *AFF4* promoter in patient-derived DMG cultures (identical reads and scale for each sample tract). **C.** AFF4 mRNA transcripts (left) and protein expression (right) in H3F3A wild-type HSJD-GBM01 and normal human astrocytes following lentiviral transduction of the H3K27M transgene. Vertical lines in indicate additional lanes removed for clarity. **D.** Genome-wide occupancy of H3K27me3 and H3K27ac before and after wild-type H3F3A re-expression in H3K27M mutant cells. **E.** Average distribution of H3K27me3 and H3K27ac peaks surrounding the transcription start site (TSS). **F.** *AFF4* promotor region before and after wild-type H3F3A re-expression demonstrates partial restoration of H3K27me3 peak, decrease in H3K27ac residue, and decrease in transcript by RNA-seq. **G.** Decrease in AFF4 mRNA transcripts (left) and protein (right) following wild-type H3F3A re-expression. **H.** Half-maximal inhibitory concentrations of atuveciclib across H3K27M+ DMG cultures, H3F3A WT pediatric glioblastoma (GBM01), immortalized astrocytes (NHA), fibroblasts (NIH3T3), and epithelial cells (RPE-NEO). **I.** Half-maximal inhibitory concentrations of atuveciclib in immortalized human astrocytes before and after H3K27M expression.

### The H3K27M mutation perturbs epigenetic regulation of AFF4

We then asked why normal developmental regulation by the SEC might be perturbed within the context of the H3K27M mutation. An interplay of activating and repressive histone post-translational modifications contributes to the net epigenetic signature and final transcriptional output from a given locus. H3K27M-mediated loss of the repressive H3K27-trimethyl mark is frequently accompanied by reciprocal gains of the activating H3K27-acetylation residue as a driver of gene expression changes in H3K27M+ DMG (*19, 43, 44*). Isogenic models examining H3K27M-mediated H3K27ac changes have shown that the majority of this deposition is restricted to cell state and lineage genes rather than affecting de novo genomic elements(*44*). To begin examining the role that H3K27 modifications might play in normal AFF4 regulation, we first analyzed reference chromatin-immunoprecipitation sequencing (ChIP-seq) datasets from neuroepithelial stem cells (ENCODE Project ENCSR978RSW), hPSC-derived oligodendrocyte precursor cells (iOPCs)(*45*), and surgical samplings of mature brain from various anatomical regions (ENCODE Project ENCSR107PPJ, ENCSR374BLY, ENCSR528DQE, ENCSR234PGX)(*46*). In either the neuroepithelial stem cell or iOPC state, the *AFF4* promoter bears an activating signature with broad H3K27ac deposition. In contrast, these H3K27ac peaks are markedly decreased across differentiated brain tissues (**Fig 4A**), suggesting a role for measured epigenetic downregulation during normal CNS development. When compared to the pattern of deposition observed in multiple patient-derived H3K27M+ DMG samples, the *AFF4* promotor consistently shows broad H3K27ac occupancy, partially recapitulating the activating precursor cell signature (**Fig 4B**). We then tested whether this predicted pattern of H3K27M-mediated AFF4 expression could be replicated experimentally. We utilized a lentiviral vector to introduce the H3.3K27M oncohistone into both H3 wild-type pediatric glioblastoma cells (HSJD-GBM001) and immortalized normal human astrocytes. In each case, the expected histone post-translational modifications were accompanied by a corresponding increase in AFF4 mRNA and protein product (**Fig 4C**).

In order to more granularly characterize the effect of the H3K27M mutation on AFF4 epigenetic repression specifically within DMG, we next created an isogenic experimental model by re-expressing a wild-type *H3F3A* transgene in H3K27M-mutant HSJD-DIPG007 cells. While not completely excising the dominant-negative effect of the H3K27M mutation, this resulted in a transient window of a genome-wide restoration of H3K27me3 and decrease in H3K27ac at transcriptional start sites as measured by ChIP-seq (**Fig 4D-E**). Partial restoration of H3K27me3 marks at the *AFF4* gene locus was accompanied by a significant concordant loss of H3K27ac and a decrease in AFF4 mRNA transcripts and protein expression (**Fig 4F-G**), consistent with reversion to a histone-modification-mediated repressed state.

Finally, we then examined sensitivity to CDK9 inhibition across a range of H3K27M-mutant DMG, H3 wild-type GBM, immortalized astrocyte, and non-glial cell cultures, we observed a gradation of CDK9i response (**Fig 4H**). Normal astrocytes were modestly sensitized to CDK9i therapy following expression of the H3K27M mutation, consistent with an oncohistone-mediated effect (**Fig 4G**). Taken together, these data support a model in which the H3K27M mutation perturbs the epigenetic regulation *AFF4* undergoes across neuroglial development and instead may allow the aberrantly-expressed SEC to contribute to DMG stem-like state maintenance. The presence of the H3K27M mutation exaggerates, but is not required for, response to CDK9 pharmacologic inhibition, suggesting this may reflect a lineage or cell state dependency, similar to observations from other epigenetic agents with therapeutic promise in DMG (*44, 47*).

### Permissive CDK9 mediation of transcriptional elongation facilitates cell growth and impedes terminal morphogenesis

Given the mechanistic role of CDK9 in Pol II pausing release, we next examined the consequences of CDK9i treatment on processive transcriptional elongation. Productive transcription by RNA Pol II requires stepwise regulatory inputs. An initiation signal is first provided by CDK7 phosphorylation of Pol II CTD at the serine 5 position (Ser5). An elongation signal is then provided by CDK9 mediated phosphorylation at serine 2 (Ser2) (*48*). Following CDK9 inhibition by atuveciclib treatment, we observed a dose-dependent abrogation of the phospho-Ser2 elongation signal with a relative preservation of the phospho-Ser5 signal for transcriptional initiation (**Fig 6A**). Together, these suggest that CDK9 inhibition by atuveciclib results in a highly selective targeting of RNA Pol II pausing release without a broader disruption of transcription initiation.

**Figure 6.**
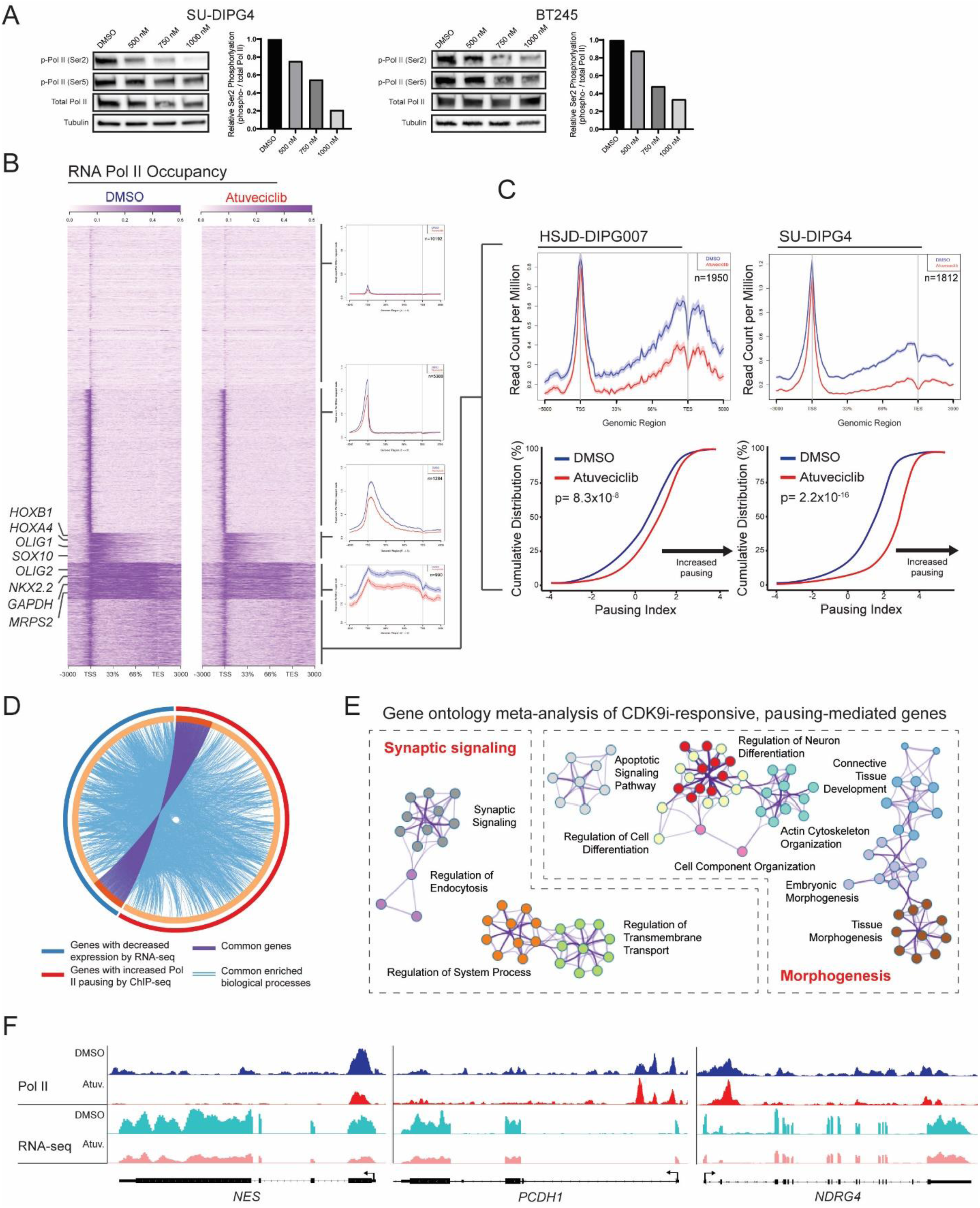
Permissive CDK9 mediation of transcriptional elongation impairs normal morphogenesis in DMG. **A.** Immunoblot of RNA Pol II CTD phosphorylation at the serine 2 (Ser2) and serine 5 (Ser5) positions following increasing doses of atuveciclib. Quantification (right) of pausing release at increasing doses of atuveciclib as measured by the ratio of Pol II phosphorylation at Ser2 relative to total Pol II. **B.** Heatmap of Pol II ChIP-seq in HSJD-DIPG007 clustered by distribution relative to transcriptional start site (TSS). Average distribution across gene body shown for each cluster. **C.** Average distribution and empirical cumulative distribution function (ECDF) plot of pausing indices for CDK9i-responsive clusters show significant increase in promoter-proximal pausing following atuveciclib treatment (two-sided Kolmogorov-Smirnov test, HSJD-DIPG007 p=8.3×10^-8^ and SU-DIPG4 p=2.2×10^-16^). **C.** Circos plot of gene ontology meta-analysis of genes with decreased expression by RNA-seq or increased Pol II pausing by ChIP-seq in HSJD-DIPG007. Purple connections indicate shared genes in both lists. Blue connections indicated common enriched biological processes. **D.** Meta-analysis reveals enrichment in networks mediating synaptic signaling and glioneuronal morphogenesis. Each node denotes an enriched term, and node color indicates enrichment contribution from respective gene list. **E.** Illustrative promoters at *NES*, *PCDH1*, and *NDRG4* loci demonstrate increased RNA Pol II pausing and decreased mRNA transcript following atuveciclib treatment.

To assess the impact of CDK9 inhibition on Pol II distribution across the chromatin landscape, we then performed ChIP-seq using Pol II antibody in DMSO- or atuveciclib-treated HSJD-DIPG007 and SU-DIPG4 cells. Consistent with whole-cell immunoblots of Pol II phosphorylation, while a modest decrease in global Pol II occupancy was observed, more than 81% and 74% of peaks present in the control state were preserved following atuveciclib treatment within a respective cell line. Unsupervised clustering of genes by distribution of Pol II occupancy revealed a hierarchy in the transcriptional architecture of DMG while clearly identifying a subset of CDK9i-responsive, pausing-mediated genes (**Fig 6B**, **Fig S3A**, and **Table S3A**). The majority of the genome displayed either an absence of Pol II (10,192 or 12,201 genes, HSJD-DIPG007 and SU-DIPG4 respectively) or detectable Pol II loading without evidence of downstream elongation (5,388 or 4,251 genes). The remaining genes with active Pol II procession into the gene body reflected both genes involved in essential cellular processes as well as a neuroglial cell lineage, but these clustered into three broad categories. The cluster (990 or 538 genes) characterized by the greatest Pol II occupancy distributed broadly across the gene body was enriched for both housekeeping genes (*GAPDH*, mitochondrial and ribosomal proteins) as well as markers of oligodendrocyte-lineage commitment (*OLIG2, NKX2.2*)(**Table S3B**) and was largely unaffected by atuveciclib treatment. A second cluster (1,284 or 1,001 genes) exhibited engaged Pol II elongation without as markedly constitutive deposition. This cluster contained genes evidencing further lineage specification (*OLIG1, SOX10*) as well as a strong anterior-posterior regionalization signature (*ALX3*, *HOXA4*, *HOXB1*, and other HOX genes)(**Table S3C**), and it exhibited a comparatively modest shift towards increased promoter-proximal pausing following atuveciclib exposure. The final cluster of 1,950 (or 1,812) genes, however, showed a marked reduction in Pol II egress into the gene body in comparison to normalized loading at the TSS. This transcriptional pausing may be quantified by defining a pausing index as a ratio of Pol II occupancy within the promoter-proximal region as compared to the gene body (*49*). Empirical cumulative distribution function (ECDF) analysis of pausing indices within this cluster of genes confirmed a significant shift towards increased promoter-proximal pausing after atuveciclib treatment in both cell lines (**Fig 6C**).

We then examined this cluster of CDK9i-responsive, pausing-mediated genes in comparison to expression changes observed by RNA-seq. Gene ontology meta-analysis delineated a subset of shared pausing-mediated genes at the core of common enriched biologic processes (**Fig 6C**, **Fig S3B**, and **Table S3D**). These processes reflected a network of gene programs with significant enrichment for neuroglial morphogenesis, with highest-ranked common enrichment in terms associated with synaptic signaling, regulation of synaptic plasticity, and the cellular morphogenesis involved in neuron projection and axonogenesis (**Fig 6D-E**, **Fig S3C**, and **Table S3E**). Together, these data affirm that the CDK9-mediated permissive regulation of transcriptional elongation at these genes contributes to the differentiation arrest observed in DMG and support the possibility that this might be reversed through CDK9i treatment.

### CDK9 inhibition can be employed for therapeutic effect in orthotopic xenograft models of DMG

Finally, we sought to determine whether this transcriptional dependency might be exploited for therapeutic effect *in vivo*. BT245 cultures were transduced with a luciferase-expressing construct and injected stereotactically into athymic nude mouse pons, generating orthotopic xenografts which anatomically and histopathologically model DIPG (**Fig 7A**). Mice were randomized sequentially 6-8 days following tumor cell implantation to either CDK9i treatment or vehicle control. An initial xenograft cohort receiving atuveciclib therapy (30 mg/kg/dose OG, three daily doses followed by a two-day break) showed only modest survival benefit in comparison to those receiving vehicle control (median survival increase of 2 days or 5%, log-rank *p=0.043). However, a second cohort treated with AZD4573 (20 mg/kg IP, three times weekly) exhibited a more dramatic benefit to overall survival (median survival increase of 8.5 days or 25%, log-rank *p=0.034) (**Fig 7B-C**), perhaps suggestive of a greater translational potential for this agent. A patient-analogous response characterization of murine xenografts using magnetic resonance imaging (MRI) with volumetric analysis was performed and showed decreased tumor growth, reduced peritumoral edema, and less invasion of surrounding anatomic structures in mice receiving CDK9i therapy (**Fig 7D** and **Fig S4**). CDK9i-treated mice experienced lymphopenia, but no significant weight loss or other treatment-associated toxicities were observed (**Fig S5**). Histologically normal-appearing neurons were observed throughout the remainder of the pons, with no evidence of treatment-related effect on uninvolved pontine tissue.

**Figure 7.**
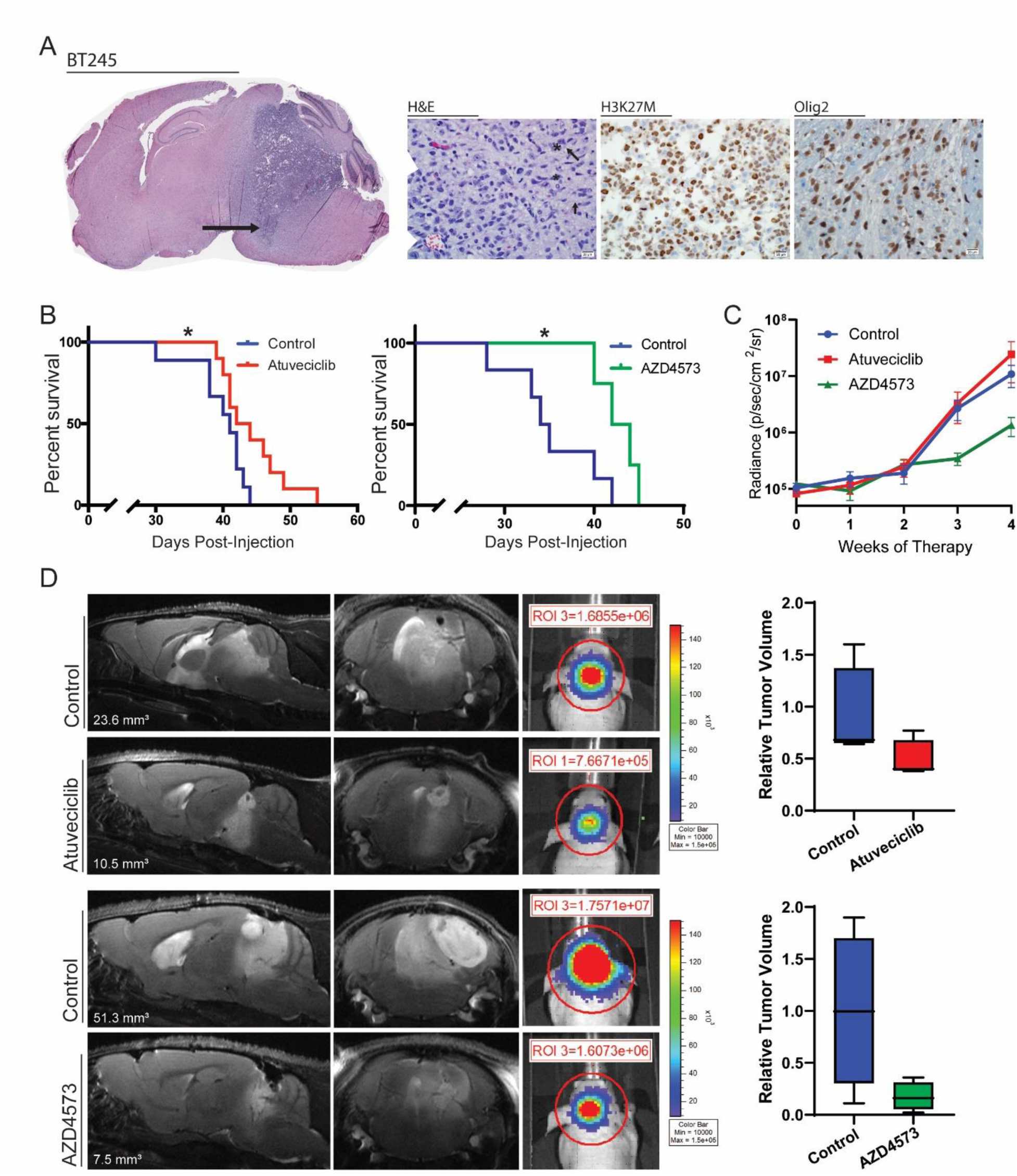
CDK9i therapy demonstrates anti-tumor effect in orthotopic xenograft models of DMG. **A.** Whole-brain sagittal H&E staining of engrafted BT245 cells shows diffusely infiltrative tumor centered in the pons (arrow) with invasion into the midbrain and thalamus. Arrows in high-power field indicate normal pontine axons surrounded by infiltrating glioma cells (asterisks). Immunohistochemical staining demonstrates H3K27M, Olig2-positive infiltrating tumor cells with normal endothelial cells, neurons, and glia seen in the background. **B.** Survival analysis of atuveciclib- (median 43 days vs 41 days, log-rank test *p=0.043, n=9 and 10, respectively) and AZD4573-treated cohorts vs vehicle control (median 43 days vs 34.5 days, log-rank test *p=0.034, n=4 and 6, respectively). **C.** Bioluminescent flux values from vehicle control, atuveciclib, and AZD4573-treated cohorts. Data are displayed as mean ± SEM. **D.** Sagittal (left) and axial (middle) T2-weighted turboRARE MRI sequences with concurrent bioluminescent imaging (right) of selected mice from vehicle control (top) and CDK9i-treated (bottom) cohorts. White text insert represents tumor volume at given timepoint. MRI volumetric analysis (right) demonstrates trend towards decreased tumor volume in each CDK9i-treated cohort (n=3 each, atuveciclib v control one-tailed t-test p=0.15, AZD4573 v control 0.056).

## Discussion

Decades of clinical trials have failed to improve outcomes or identify effective medical therapies for children diagnosed with DMG (*2, 3*). Insights stemming from the identification of the oncohistone H3K27M as the unifying driver of these tumors, however, have introduced the potential for therapies targeting the dysregulated epigenetic milieu imparted by these recurrent mutations. Recognition of the aberrancies introduced across the epigenetic regulatory landscape by the H3K27M mutation has led several groups to explore and define epigenetic therapies such as histone demethylase inhibition (*50*) or histone deacetylase inhibition (*20, 47, 51, 52*) as promising therapeutic strategies for DMG. Several of these agents are now moving forward in phase I clinical trials (NCT02717455, NCT03632317, NCT03566199) but have yet to demonstrate clinical efficacy. As such, there exists growing interest in alternative means of targeting the H3K27M oncohistone, including the proximate transcriptional consequences of this mutation. Our data establishes that epigenome reprogramming by H3K27M aberrantly activates machinery involved the regulation of transcriptional elongation, identifying a consequent dependency that can be targeted for therapeutic effect.

Transcriptional regulation is a complex synthesis of numerous oppositional or enhancing regulatory inputs. Many developmentally-regulated genes are marked by paused RNA Pol II within their promotor-proximal regions (*28, 34, 35, 49, 53, 54*), which may allow for the synchronous induction of gene programs within a distinct cell population (*28, 53*). In normal cellular contexts, the super elongation complex incorporates the CDK9-containing P-TEFb from an inactive, sequestered pool in order to promote rapid transcriptional induction (*24, 26*). The SEC was originally characterized as a consequence of the frequent in-frame MLL*-*SEC-member fusions identified in *MLL-*rearranged childhood leukemias, where it was found to mediate the transcriptional changes observed in that disease model (*22, 23*). Misregulation of transcriptional elongation checkpoint control has since been implicated in several malignancies and other human diseases (*23–25, 29*). In DMG, the disordered expression of SEC may disrupt the carefully orchestrated, context-dependent activation under which pausing-regulated expression programs ought to normally proceed.

As a result of the growing clinical interest in more nuanced therapeutic manipulation of cellular transcription, multiple CDK9 small-molecule pharmacologic inhibitors have entered the drug development pipeline (*30, 39*). Unlike the chemical tools often used to study transcription, many of these are clinically relevant compounds. To date, at least six CDK9 inhibitors have demonstrated sufficient antineoplastic activity *in vitro* to be carried forward in phase I or II clinical trials (*39, 55–59*). Of the newer generation, highly-selective inhibitors, atuveciclib and AZD4573 are both advancing in clinical development, with adult phase I trials currently in progress (NCT01938638, NCT02345382, NCT03263637). The efficacy of AZD4573 demonstrated in this study as well as the growing clinical experience with the dosing and safety profiles of this class of medications positions this agent as ideally suited for rapid translation to pediatric trials.

In summary, we have used a targeted, unbiased screening approach to epigenetic regulation in H3K27M-mutant DMG to identify the SEC as a novel, targetable dependency. Dysregulation of the SEC as a consequence of the H3K27M mutation appears to drive a permissive transcriptional program which contributes to DMG stem-cell maintenance. Inhibition of this pro-oncogenic signaling via CDK9i treatment restores promotor-proximal pausing of Pol II and permits cellular differentiation programs to proceed. The anti-tumor activity demonstrated in orthotopic xenograft models of DMG using clinically-relevant CDK9 inhibitors could prove transformative if translated into clinical trials for children diagnosed with this devastating disease.

## Materials and Methods

### Cell lines and primary patient samples

DMG cells were maintained as previously described (*60*). Briefly, SU-DIPG4 cells were provided by Dr. Michelle Monje (Stanford University, California) and cultured in tumor stem media (TSM) consisting of Neurobasal(-A) (Invitrogen), B27(-A) (Invitrogen), human-basic FGF (20 ng/mL; Shenandoah Biotech), human-EGF (20 ng/mL; Shenandoah Biotech), human PDGF-AB (20 ng/mL; Shenandoah Biotech) and heparin (10 ng/mL). HSJD-DIPG007 cells were provided by Dr. Angel Montero Carcaboso (Sant Joan de Déu, Barcelona) and maintained in TSM media as above with 10% fetal bovine serum (FBS)(Atlanta Biologicals). SF8628 cells were provided by Dr. Nalin Gupta (University of California, San Francisco) and maintained in Dulbecco’s modified eagle medium (DMEM)(Gibco/ThermoFisher) and supplemented with 10% FBS, MEM non-essential amino acids (Gibco/ThermoFisher) and antibiotic/antimicotic (Gibco/ThermoFisher). BT245 cells were provided by Dr. Adam Green (University of Colorado) and grown in NeuroCult NS-A media (Stemcell Technologies) supplemented with penicillin-streptomycin (1:100), heparin (2 μg/mL), human epidermal growth factor (EGF; 20 ng/mL), and human basic fibroblast growth factor (FGFb; 10 ng/mL). Cells were grown in either adherent monolayer conditions (Falcon/Corning) or as tumor neurospheres (ultra-low attachment flasks, Corning) as indicated. All cell lines were validated by DNA fingerprinting through the University of Colorado Molecular Biology Service Center utilizing the STR DNA Profiling PowerPlex-16 HS Kit (DC2101, Promega)(**Table S4**).

### shRNA screening

SU-DIPG4 cell line was transduced with a pooled lentiviral shRNA library consisting of ∼4200 shRNAS targeting 408 epigenetic genes (approximately 4–10 shRNAs per gene, 4188 total shRNAs). Cells were infected with the pooled shRNA lentiviral library with a multiplicity of infection (MOI) of 0.3 using polybrene on day 1. One aliquot (control) of cells was collected for isolating genomic DNA at 4 days, and the transduced cells were selected with puromycin treatment. These cells were maintained for a further 21 days (passaging every 3 days). Cells were passaged to maintain an MOI of 0.3 at all times to ensure that each transduced cell had a single genomic integration from the shRNA expressing the same gene. The cells were then collected at 9 days and finally at 21 days (approximately 6 doubling times) after transduction. Genomic DNA was isolated from these transduced cells 4 and 21 days after transduction, processed to amplify individual shRNAs with two rounds of PCR, and sequenced with an Illumina Hiseq instrument. The sequencing results were analyzed using R2 by comparing the shRNAs present on day 4 to day 21, with FDRs of 0.5 and 0.1, respectively.

### shRNA transduction

HEK293 FT cells were used to produce the viral particles by packaging the vectors, PSPAX2 and PMD2.G with the shRNA plasmids. shNull and shAFF4 plasmids were purchased from transOMIC (RLGH-GU01145 and ULTRA-3271665). shCDK9 plasmids were purchased from the University of Colorado Functional Genomics Core Facility (TRCN0000000494 and TRCN0000000498). H3K27M transgene and H3F3A-overexpression vectors were kindly provided by Dr. C. David Allis (Rockefeller University, NY). Approximately 750,000 target DMG cells were seeded onto a 10 cm2 tissue culture plate (Falcon/Corning, Durham NC) and transduced using polybrene. Virus-containing media was removed at 24 hours, and cells were selected in puromycin 1 ug/mL after 48 hours and subsequently maintained in puromycin-containing media.

### Proliferation assays

Cell proliferation assessed using xCELLigence Real-Time Cell Analysis (RTCA) machine and software. Cells (2,000 – 6,000 cell/well) were seeded onto a gold-plated 16- or 96-well E-plate, and impedance of cells are measured over time. Cell index calculations performed using RTCA software.

### Colony focus assay

Cells were treated to achieve stable shRNA transductions as described above. Then 500 cells/well were seeded on a 6-well tissue culture plate (Corning Costar/Fisher) and allowed to grow for 14 days. At that time, cells were stained with 0.25% crystal violet, and colonies containing >25 cells were manually counted under light microscopy.

### Neurosphere assay

Cells were treated to achieve stable shRNA transductions as described above. 250 cells/well were seeded on a 96-well ultra-low-attachment round-bottom tissue culture plate (Corning) and centrifuged to promote neurosphere formation. Plates were then monitored on IncuCyte S3 Live Cell Analysis System for 14 days. Measurements of neurosphere size over time using GFP reporter were calculated by the IncuCyte S3 software and normalized against day 1 values to determine growth.

### Aldehyde dehydrogenase assay

The ALDH activity of was measured using Aldefluor kit (Stem Cell Technologies) according to the manufacturer’s instruction. Briefly, 1 × 10^5^ cells were resuspended in 0.5 ml Aldefluor buffer, separated equally into two tubes, one of which was added 5 ul of DEAB reagent as negative control. Then 1.25 ul of Aldefluor Reagent was added to each tube and mixed well. After incubation at 37°C for 45 min and centrifugation, cells were stained with propidium iodide and then analyzed on the Guava easyCyte HT flow cytometer (Luminex).

### Extreme limiting dilution assay (ELDA)

For shRNA studies, cells were treated to achieve stable shRNA transductions as described above. For drug treatment studies, cells were treated with indicated concentration prior to re-plating. Limiting dilution assay was then performed as previously described(*61*). Briefly, cells were seeded on a 96-well ultra-low-attachment round-bottom tissue culture plate (Corning) in serum-free media at increasing numbers from 1 cell/well to 250 cells/well. Cells were seeded from n=5 wells (250 cells/well, 100 cells/well), n=10 wells (10-50 cells/well), or n=30 wells (1 cell/well) per condition. Cells were allowed to grow for 14 days, and the number of wells containing neurospheres was counted manually under light microscopy. Published ELDA software (http://bioinf.wehi.edu.au/software/elda/) was used to calculate comparative self-renewal potential of cells.

### Co-Immunoprecipitation

Immunoprecipitation experiments were performed with whole cell extracts. Cell lysates were immunoprecipitated using Universal Magnetic Co-IP Kit (Active Motif 54002), then analyzed by western blot. Immunoprecipitation was performed using anti-AFF4 antibody and normal rabbit IgG (see Key Resources Table). Antibody/extract mixtures were incubated with complete Co-IP/wash buffer on a 4 degree shaker for 4 hours. Magnetic protein G beads were added and incubated another hour, then washed using the complete co-IP/wash buffer and a magnetic stand. Bead pellets were re-suspended in 1X reducing loading buffer for immunoblotting.

### Drug screening

The small molecule inhibitors PHA-767491, flavopiridol, and LDC000067 were obtained from SelleckChem. The small molecule inhibitors atuveciclib (BAY-1143572) and AZD4573 were obtained from MedChemExpress. All drugs were dissolved in DMSO per vendor instructions to make stock concentrations for use in *in vitro* experiments. 5000 cells were seeded per well (n=4) in a 96 well plate, and the corresponding drug or DMSO control was added 24 hours later. Cells were allowed to grow for 5 days before adding an MTS reagent (Promega) and measuring absorbance at 492 nM on a BioTek Synergy H1 microplate reader (BioTek Instruments, VT). A ratio of absorbance in treated vs DMSO samples was used to generate a heatmap of screening results.

### Transcriptome sequencing (RNA-seq)

Ribonucleic acid was isolated from cells in indicated experimental conditions using a Qiagen miRNAeasy kit (Valencia, CA) and measured on an Agilent Bioanalyzer (Agilent Technologies). Illumina Novaseq 6000 libraries were prepared and sequenced by Novogene (CA, USA) or Genomics and Microarray Core Facility at the University of Colorado Anschutz Medical Campus. High-quality base calls at Q30 ≥ 80% were obtained with ∼40 M paired-end reads. Sequenced 150bp pair-end reads were mapped to the human genome (GRCh38) by STAR 2.4.0.1, read counts were calculated by R Bioconductor package GenomicAlignments 1.18.1, and differential expression was analyzed with DESeq2 1.22.2 in R. Further analysis by GSEA was performed in GSEA v2.1.0 software with 1,000 data permutations.

### Chromatin immunoprecipitation sequencing (ChIP-seq)

The growth medium was aspirated from cells (70% confluence) and replaced with a 1% formaldehyde in 1X DPBS for 9 min at room temperature. The cross-linking was terminated by addition of glycine to a final concentration of 0.125 M and incubation for 5 min. The cells were washed twice with cold PBS and scraped in ChIP lysis buffer (1% SDS, 10 mM EDTA and 50 mM Tris-HCl pH 8.1) containing COmplete EDTA free protease inhibitors (Sigma). The cell lysates were sonicated with a Bioruptor Plus (Diagenode) for 25 cycles (30 seconds ON and 30 seconds OFF). The size of the sonicated DNA fragments was checked on an 1X TAE1 % agarose gel electrophoresis and ranged between 250 to 500 bp. After removing the debris via centrifugation, chromatin extracts were collected and protein concentration was determined by BCA protein assay reagents (Pierce).

For immunoprecipitation, chromatin extracts with 1 mg of total protein for each sample were incubated with primary antibodies (dilution 1:50) overnight at 4°C. Primary antibodies used are included in table under “Western blotting” below. After incubation with the primary antibody, 20 uL of pre-washed magnetic beads (Magna ChIP Protein A+G Magnetic Beads, Millipore Sigma) were added to each sample for 2 hours at 4°C. Using a magnetic rack, the immunoprecipitates were washed successively with 1 ml of low salt buffer (20 mM Tris-HCl [pH 8.0], 150 mM NaCl, 0.1% SDS, 1% triton X-100, 2 mM EDTA), high salt buffer (20 mM Tris-HCl [pH 8.0], 500 mM NaCl, 0.1% SDS, 1% triton X-100, 2 mM EDTA), LiCl washing buffer (10 mM Tris-HCl [pH 8.0], 250 mM LiCl, 1.0% NP40, 1.0% deoxycholate, 1 mM EDTA) and twice with TE buffer. The DNA-protein complexes were eluted with 300 µl of IP elution buffer (1% SDS, 0.1 M NaHCO3). The eluents were pooled together and the cross-links were reversed by adding NaCl (a final concentration 0.2 M) into the eluents and incubating them at 65 °C overnight. The DNAs were recovered by proteinase K and RNase A digestion, followed by phenol/chloroform extraction and ethanol precipitation. Pellets were resuspended in 50 µl of 10 mM Tris-HCl [pH 8.0]. ChIP-DNA was quantified using the Qubit dsDNA High Sensitivity Assay kit (Thermo Fisher Scientific).

ChIP-seq libraries were sequenced on the Illumina NovaSEQ6000 platform. To remove Illumina adapters and quality-trim the read ends, reads were filtered using BBDuk (http://jgi.doe.gov/data-and-tools/bb-tools<u>). Bowtie2 was used to align the 150-bp paired-end sequencing reads to the hg38</u> reference human genome. Samtools (v.1.5) was used to remove unmapped mapped reads and to randomly extract the same number of reads for all samples. Peaks were called using MACS2 (v2.1.1.20160309)(*62*) with default parameters. Peak locations were further annotated using the ChIPseeker R package (*63*). For pausing indices, the promoter region was defined as -200 bp upstream to +400 bp downstream and the body region was the remainder of the entire gene body. The pausing index was defined as the ratio of the average coverage of the promoter over the average coverage of the gene body. The average coverage of each promoter and each gene body was calculated with the valr R package (https://cloud.r-project.org/package=valr) using the bed_coverage function. ECDF plots were made in R version 3.5.0 using the ecdf function.

### Quantitative polymerase chain reaction

Ribonucleic acid was isolated from cells in indicated conditions using a Qiagen miRNAeasy kit (Valencia, CA). TaqMan gene expression primers and proves for *AFF4* (Hs00969465_m1), *CDK9* (Hs00977896_g1), and *GAPDH* (Hs02758991_g1) were purchased from Applied Biosystems (Carlsbad, CA). Assays were performed in triplicate according to manufacturer recommendations. GAPDH was used as an endogenous control and the gene expression relative quantity was calculated using the ΔΔCt method. Gene expression assays were performed on an ABI StepOnePlus Real-Time PCR system.

### Western blotting

Whole-cell protein lysates were harvested in lysis buffer (RIPA buffer supplemented with protease inhibitor (Roche), sodium vanadate and sodium molybdate) from cells in indicated experimental conditions. Histone proteins were isolated by acid extraction from cells. Protein concentrations were determined by BCA assay, and 1 - 3 µg of histone proteins and ∼25-50 µg of non-histone proteins were used for western blotting. Protein was separated on a 4-20% PROTEAN TGX Gels (Biorad) and blotted using a wet transfer system (Biorad). See Key Resources table for antibodies utilized.

### Orthotopic xenograft experiments

Athymic nude mice aged 4 to 8 weeks were anesthetized with 4% and maintained with 1 to 3% isoflurane (1 L/min oxygen flow). Mice were immobilized on a Kopf Model 940 Stereotaxic Frame with a Model 923 mouse gas anesthesia head holder and Kopf 940 Digital Display. Artificial Tears ophthalmic ointment was applied to the eyes. A linear incision was made to expose the skull and 30% hydrogen peroxide used to identify lambda. To target the pons, a 1.0 mm diameter burr was drilled in the cranium using a Dremel drill outfitted with a dental drill bit at 1.000 mm to the right and 0.800 mm posterior to lambda. A suspension of BT245 cells was prepared by suspending in NeuroCult NS-A serum-free media as described above to a concentration of 100,000 cells/2 µl/injection and slowly injected 5.000 mm ventral to the surface of the skull using an UltraMicroPump III and a Micro4 Controller (World Precision Instruments) equipped with a Hamilton syringe and a 26 gauge needle. The burr hole was sealed with bone wax and 1 mg/kg bupivacaine applied. The incision closed with 5-0 poly vicryl sutures (Ethicon) and a topical antibiotic applied. Post-surgical pain was controlled with SQ carprofen (5 mg/kg/daily) on the day of and for 2 days following the procedure.

Mice were randomized sequentially at 6-8 days following injection. Atuveciclib was suspended in DMSO per vendor recommendations. Treatment dosages were prepared with 80% polyethylene glycol 400 (Millipore), 10% sterile water, and 10% DMSO stock. Mice were treated with 30 mg/kg/dose via oral gavage daily for three days followed by two days break per cycle. Mice were treated with a total of 9 cycles or until they reached protocol endpoint. AZD4573 was prepared to 10% DMSO stock and diluted in 40% sterile water, 39% polyethylene glycol 400, and 1% Tween 80. Mice were treated with 20 mg/kg/dose IP three times weekly for total of 6 cycles or until reaching protocol endpoint. University of Colorado Institutional Animal Care and Use Committee (IACUC) approval was obtained and maintained throughout the conduct of the study.

### Animal imaging

All non-invasive MRI scans were performed on an ultra-high field Bruker 9.4 Tesla BioSpec MR scanner (Bruker Medical, Billerica, MA) equipped with a mouse head-array RF cryo-coil. The mouse was anesthetized with 2% isoflurane and placed on a temperature-controlled animal bed. Non-gadolinium multi-sequential MRI protocol was applied to acquire (i) high-resolution 3D T2-weighted turboRARE (**<u>R</u>**apid **<u>A</u>**cquisition with **<u>R</u>**elaxation **<u>E</u>**nhancement, 52 micron in-plane resolution); (ii) sagittal FLAIR (**<u>F</u>**luid **<u>A</u>**ttenuated **<u>I</u>**nversion **<u>R</u>**ecovery) for CSF suppression and edema; and (iii) axial fast spin echo DWI (**<u>D</u>**iffusion **<u>W</u>**eighted **<u>I</u>**maging) with six b-values for tumor cellularity and edema. All MRI acquisitions and image analysis were performed using Bruker ParaVision 360NEO software. For volumetric assessments, free-hand drawn regions of interests (ROIs) were placed over the tumor or edema region on each axial slice. The total tumor or edema volume was reported in mm3 as the sum of all ROIs from individual slices multiplied by a slice thickness (0.7 mm, no slice gap). Oval or circular ROI were used for DWI analysis and calculations of the apparent diffusion coefficients (ADCs) in the tumor. The frankly hemorrhagic foci were excluded from ADC analysis. All MRI acquisitions and image analysis was performed by a radiologist blinded to the treatment assignment of the mice.

### Immunohistochemistry

Tissue was fixed in 10% formalin, sectioned by A.G., and submitted to the University of Colorado Denver Tissue Histology Shared Resource for staining.

### Statistical analysis

Unless indicated otherwise in the figure legend, all in vitro data are presented as mean ± SEM. In vitro proliferation, clonogenic, neurosphere, qPCR, and aldehyde dehydrogenase assays, as well as drug screens and RNA sequencing were performed with a minimum of three independent samples, and key experiments were successfully replicated with independent samples on separate days. Unless otherwise specified, all statistical analysis was performed on GraphPad Prism 8.0 software (GraphPad, La Jolla, CA). For purposes of RNA-sequencing analysis, genes were considered significantly differentially expressed if meeting fold change </> 2 and p-value <0.01. Normalized enrichment scores and estimated false discovery rate as defined by GSEA are listed in figure panels. Kaplan-Meier survival curve comparisons were performed by log-rank (Mantel-Cox) test using GraphPad Prism 8.0 software.

## Acknowledgments

We would like to thank the following individuals for generously providing the cell lines used for this work: Michelle Monje (Stanford University) for SU-DIPG4 and SU-DIPG13, Angel Montero Carcaboso (Sant Joan de Déu) for HSJD-DIPG007, and Nalin Gupta (University of California, San Francisco) for SF8268.

## Author contributions

S.F. and S.V. performed the pooled shRNA screening. N.D. and F.M.W. performed the shRNA transductions and phenotypic assays. D.W. performed the RNA-sequencing alignment and analysis. N.D. and K.M. performed the qPCR experiments. N.D., F.M.W, and I.B. performed immunoblotting. F.M.W. performed the Co-IP experiments. N.D. and E.D. performed the ChIP-sequencing experiments, and E.D. was responsible for ChIP-seq alignment and analysis.

A.L.G. developed the DMG xenograft model. A.P. was responsible for the stereotactic xenograft injections; N.D. and I.B. conducted the subsequent in vivo studies. N.J.S. performed the xenograft MR imaging and was responsible for its analysis. A.G. performed neuropathologic review of brain tissue. N.K.F. procured tissue specimens and maintained IRB compliance. N.D., S.V., and R.V. contributed to data analysis and interpretation. N.D. prepared the figures and wrote the manuscript. S.V. and R.V. conceived the project, supervised all aspects of the work, and edited the manuscript.

## Data and materials availability

The accession number for the raw and processed ChIP-seq and RNA-seq data reported in this paper is GEO: GSE129777.

## Supplementary Materials

**Figure S1. Related to Figure 2.**
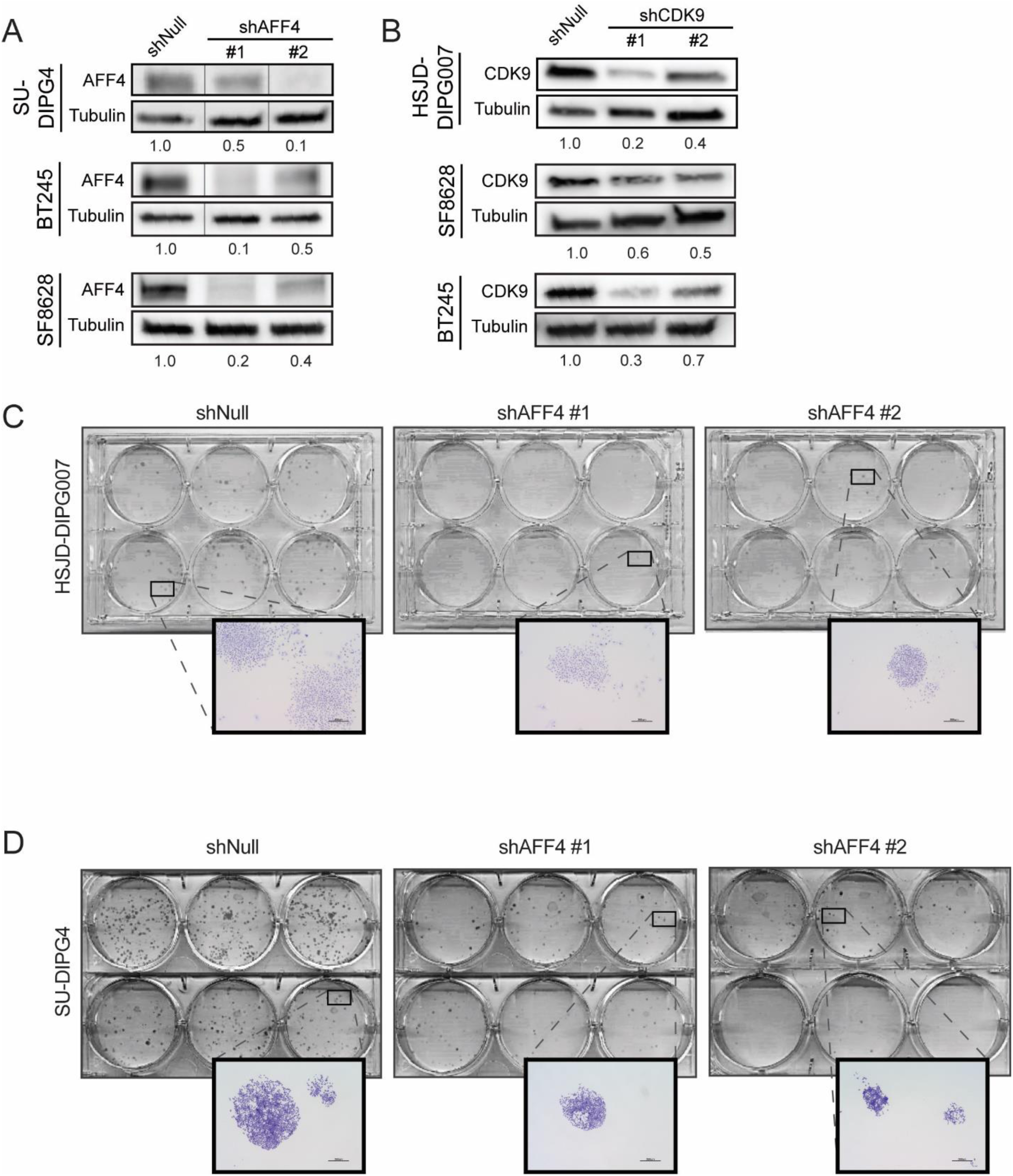
SEC member depletion diminishes clonogenic potential. Decrease in AFF4 (**A**) and CDK9 (**B**) protein in respective cell lines following transduction with two non-overlapping shRNA lentiviral constructs in comparison to shNull control. Numbers below immunoblot represent relative densitometry quantification with respect to shNull control. Vertical lines indicate additional lanes removed for clarity. **C-D.** Images of whole plate and 4x brightfield microscopy (insert) of clonogenic assays following shRNA-mediated AFF4 depletion.

**Figure S2. Related to Figure 4.**
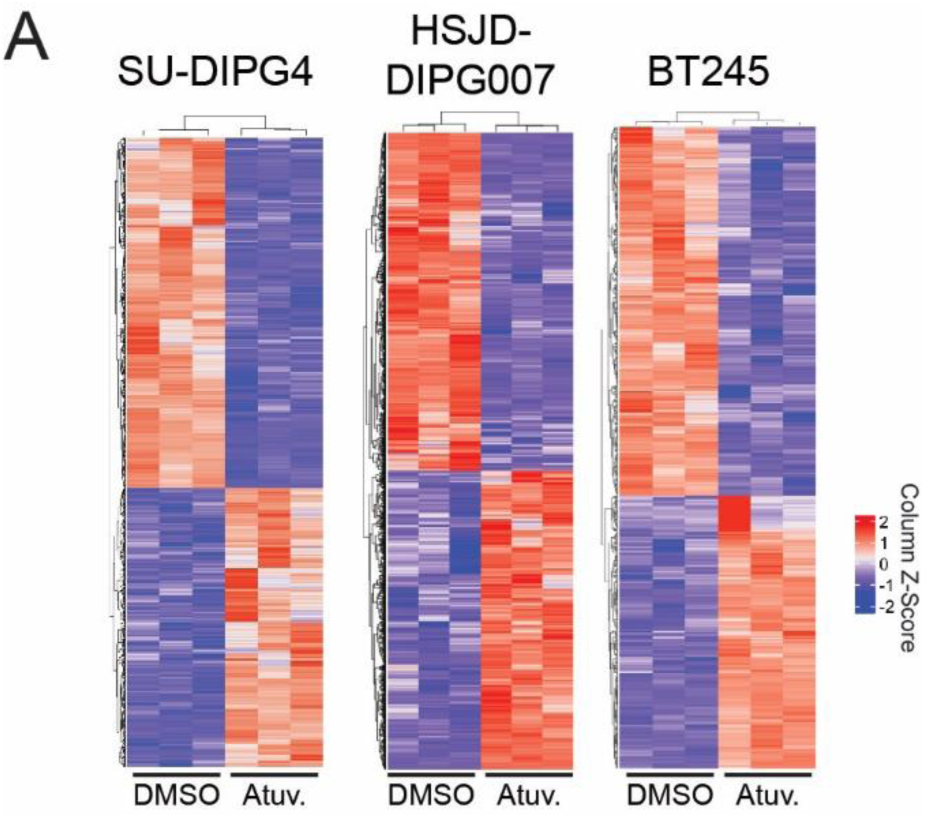
CDK9i treatment induces conserved differential gene expression across DMG cultures. Unsupervised hierarchical clustering of 1,182 differentially expressed genes from SU-DIPG4 cells (left), 1,447 genes from HSJD-DIPG007 (middle), or 918 genes from BT245 (right) treated with either atuveciclib or DMSO (n=3 each, log fold change >1 and p < 0.05).

**Figure S3. Related to Figure 6.**
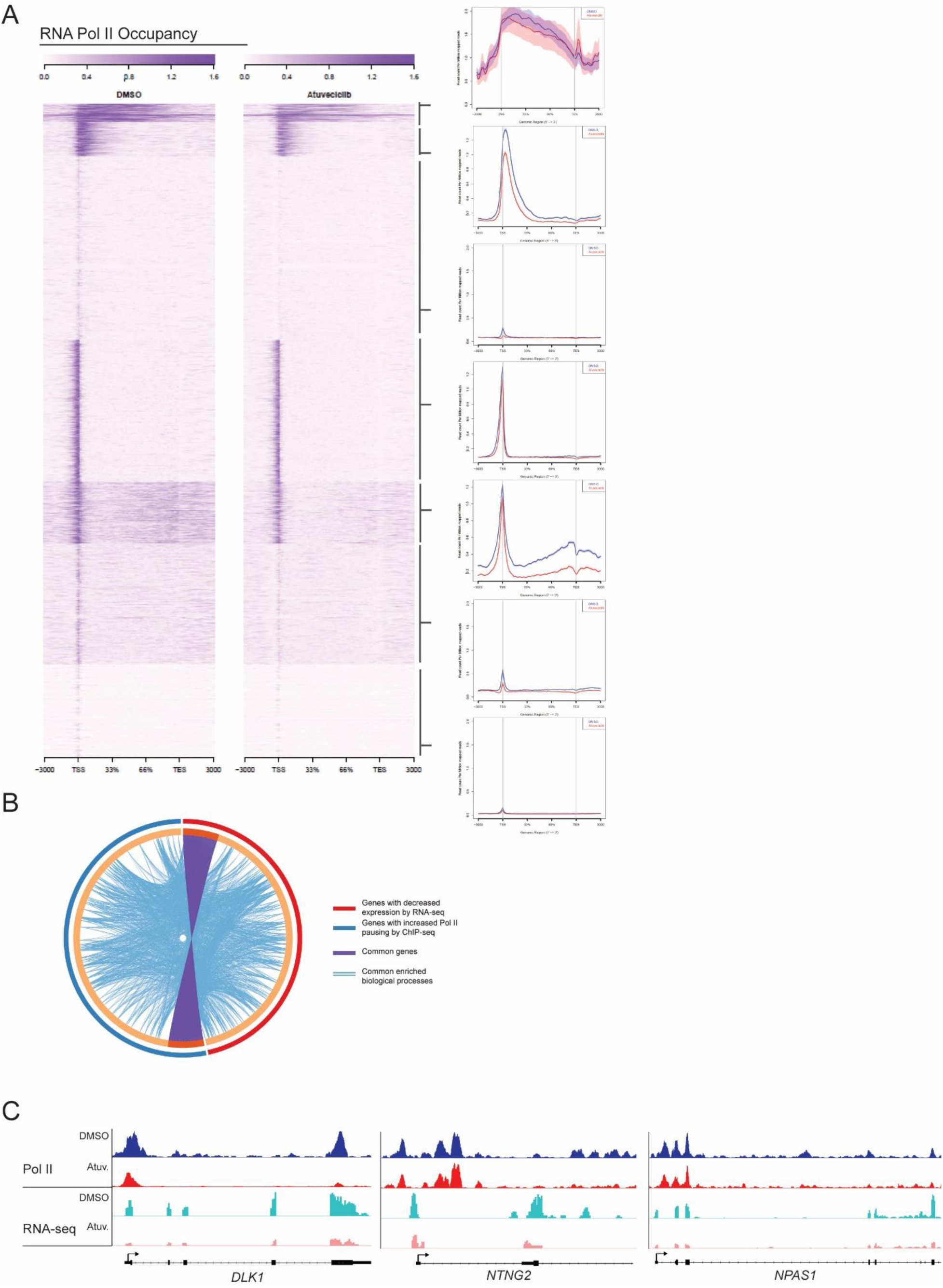
CDK9i treatment alters Pol II occupancy at defined subset of genes in DMG. **A.** Heatmap of Pol II ChIP-seq in SU-DIPG4 clustered by distribution relative to transcriptional start site (TSS) (left). Average distribution across gene body shown for each cluster (right). **B.** Circos plot of gene ontology meta-analysis of genes in SU-DIPG4 with decreased expression by RNA-seq or increased Pol II pausing by ChIP-seq. Purple connections indicate shared genes in both lists. Blue connections indicated common enriched biological processes. **C.** Illustrative promoters at *DLK1*, *NTNG2*, and *NPAS1* loci demonstrate increased RNA Pol II pausing and decreased mRNA transcript following atuveciclib treatment.

**Figure S4. Related to Figure 7.**
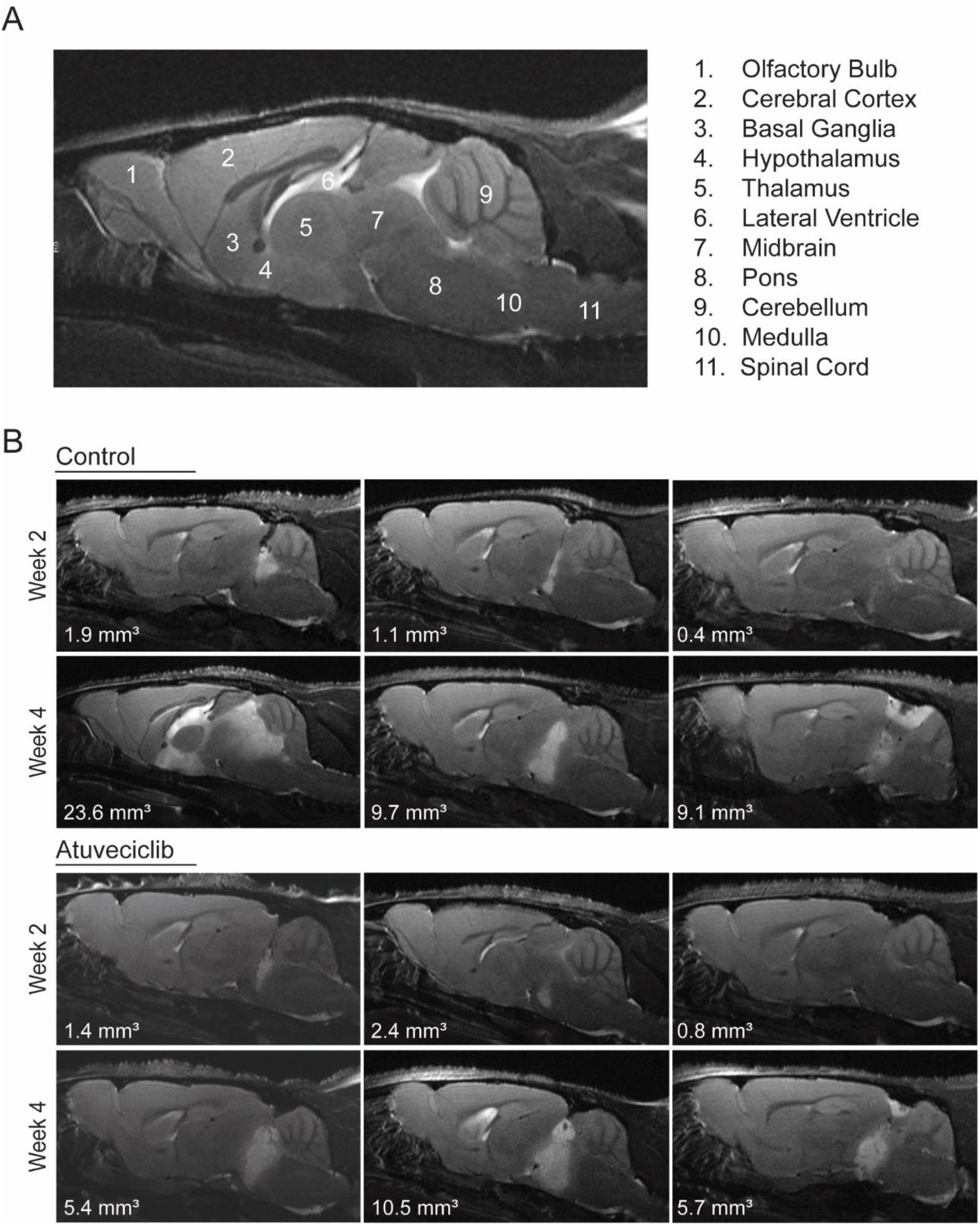
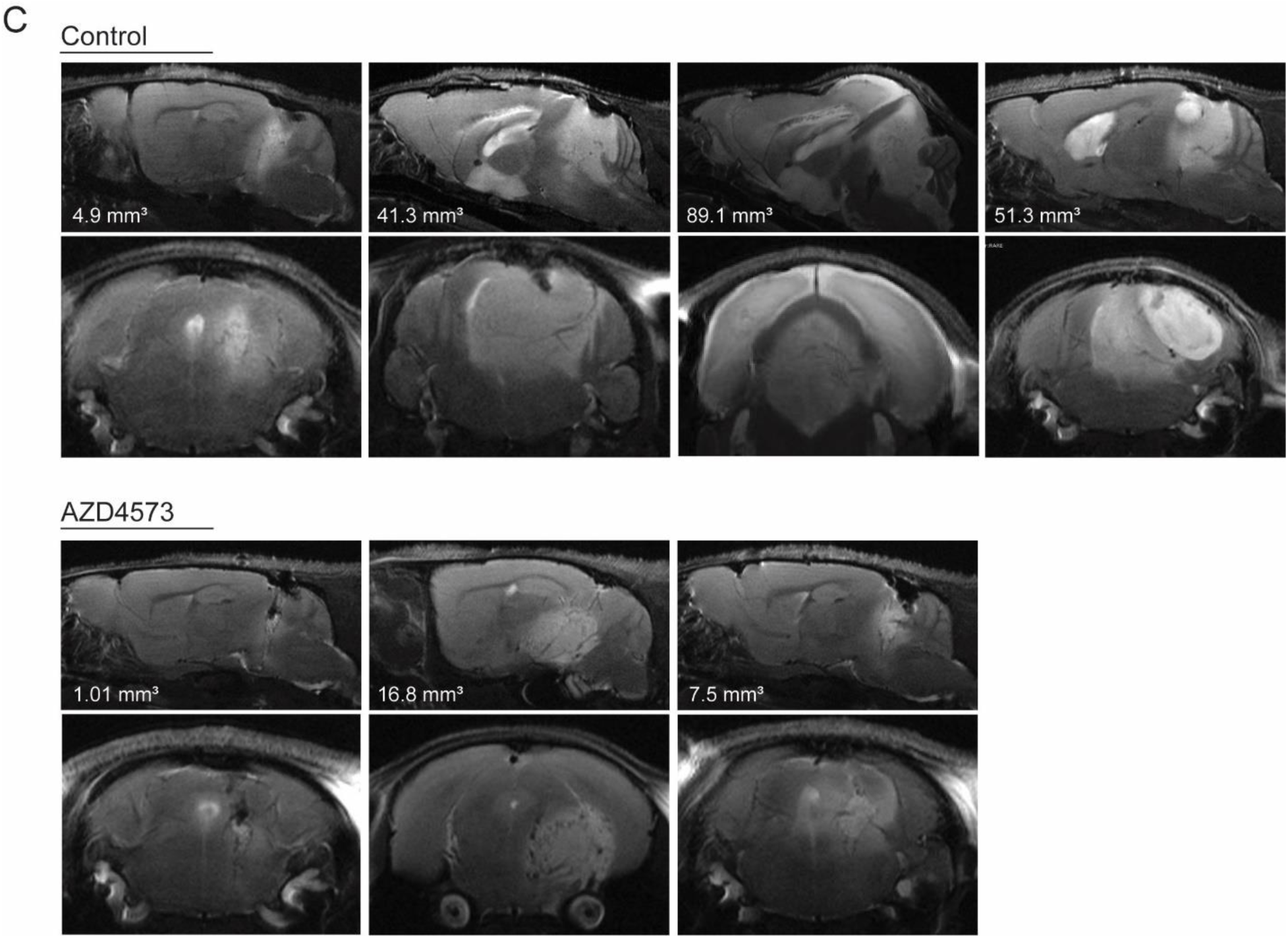
MRI demonstrates anti-tumor effect of CDK9i therapy in vivo. **A.** Sagittal T2-turboRARE sequence of normal mouse brain with key anatomical structures labeled. **B.** Sagittal T2-turboRARE sequences of select mice from vehicle control- (top) and atuveciclib-treated (bottom) mice show decreased tumor growth, peritumoral edema, and cerebellar and spinal invasion over time. White text insert represents volumetric analysis of tumor size at given time point (n=3 per group). **C.** Sagittal T2-turboRARE sequences of select mice from vehicle control- (top) and AZD4573-treated (bottom) mice demonstrate decreased tumor volume, decreased peritumoral edema, and increased necrosis. White text insert represents volumetric analysis of tumor size (n=3 per group).

**Figure S5. Related to Figure 7.**
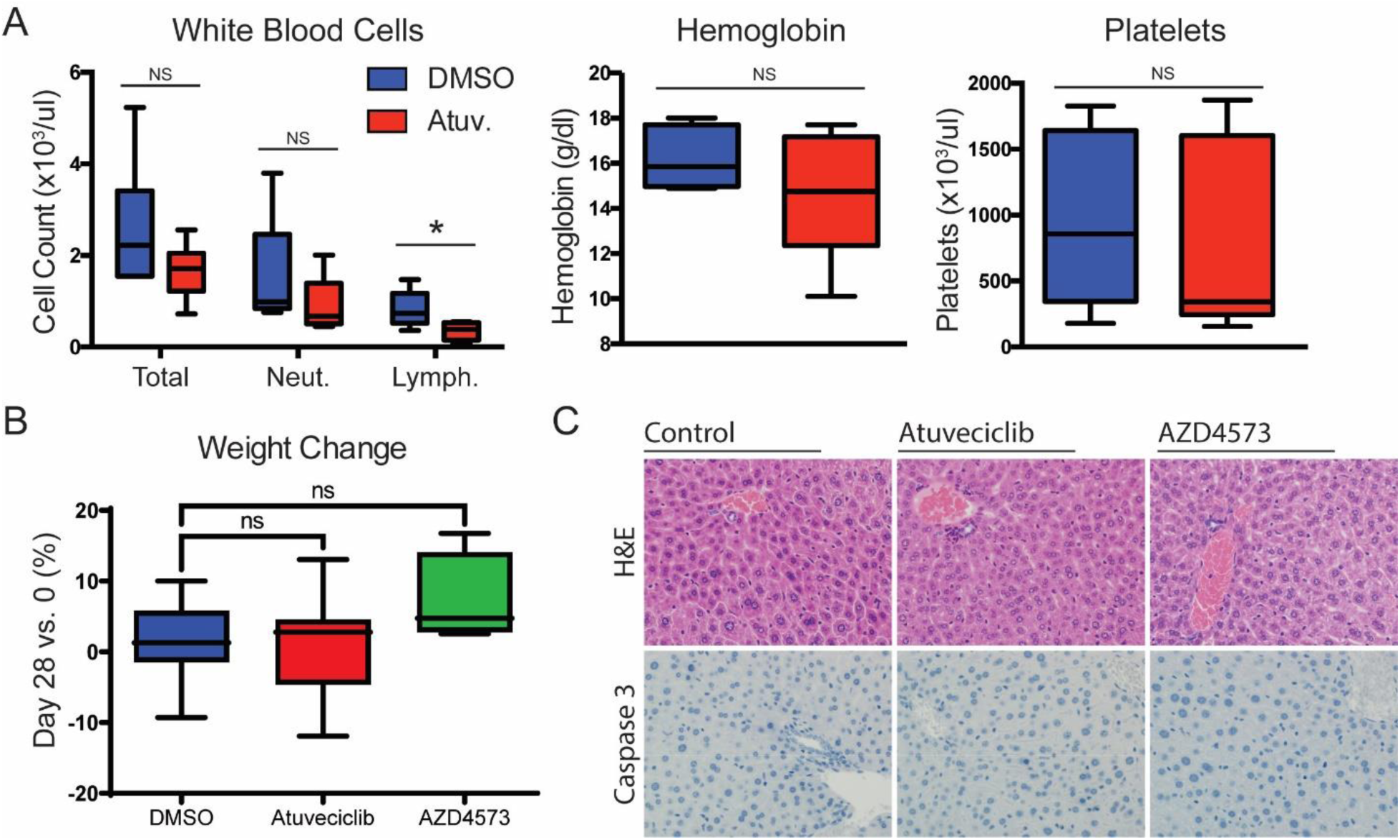
CDK9i therapy is well tolerated in murine xenograft models. **A.** Complete blood counts from sampling of vehicle control (blue) and atuveciclib treatment (red) cohorts (n=6 each). Atuveciclib treated mice show significant lymphopenia (two-tailed t test, p=0.026) and a trend towards generalized myelosuppression which did not reach statistical significance (two-tailed t test, total WBC p=0.16, neutrophils p=0.26, hemoglobin p=0.22, platelets p=0.631). **B.** Vehicle control (blue), atuveciclib treatment (red), and AZD4573 (green) cohorts show no statistically significant difference in weight change over 28 days of treatment (median +2.22% vs +2.79% vs +4.77%, two-tailed t test, p=0.58 and p=0.15, respectively). **C.** Hematoxylin and eosin (top) and caspase-3 (bottom) staining of liver sampled from vehicle control (left), atuveciclib (center), and AZD4573 treated cohorts (n=3 each). Histology demonstrates no evidence of hepatic fibrosis or caspase-mediated apoptosis suggestive of hepatocellular toxicity.

**Table S1. Related to Figure 1. Epigenomic shRNA screen in DMG.** Spreadsheet providing full list of genes included in shRNA screen, statistical cut-offs, and list and functional enrichment of hits.

**Table S2. Related to Figure 4. Gene-ontology meta-analysis of common differentially expressed genes in shAFF4 depletion and atuveciclib treatment.** Spreadsheet providing full annotation and enrichment statistics of meta-analysis comparing shAFF4 depletion and atuveciclib treatment in BT245 cells.

**Table S3. Related to Figure 6. RNA Pol II pausing and meta-analysis.** Spreadsheet containing full list of genes by Pol II clustering in HSJD-DIPG007 and SU-DIPG4 cells, as well as gene-ontology meta-analysis correlating Pol II pausing with observed RNA-seq expression changes.

**Table S4.**
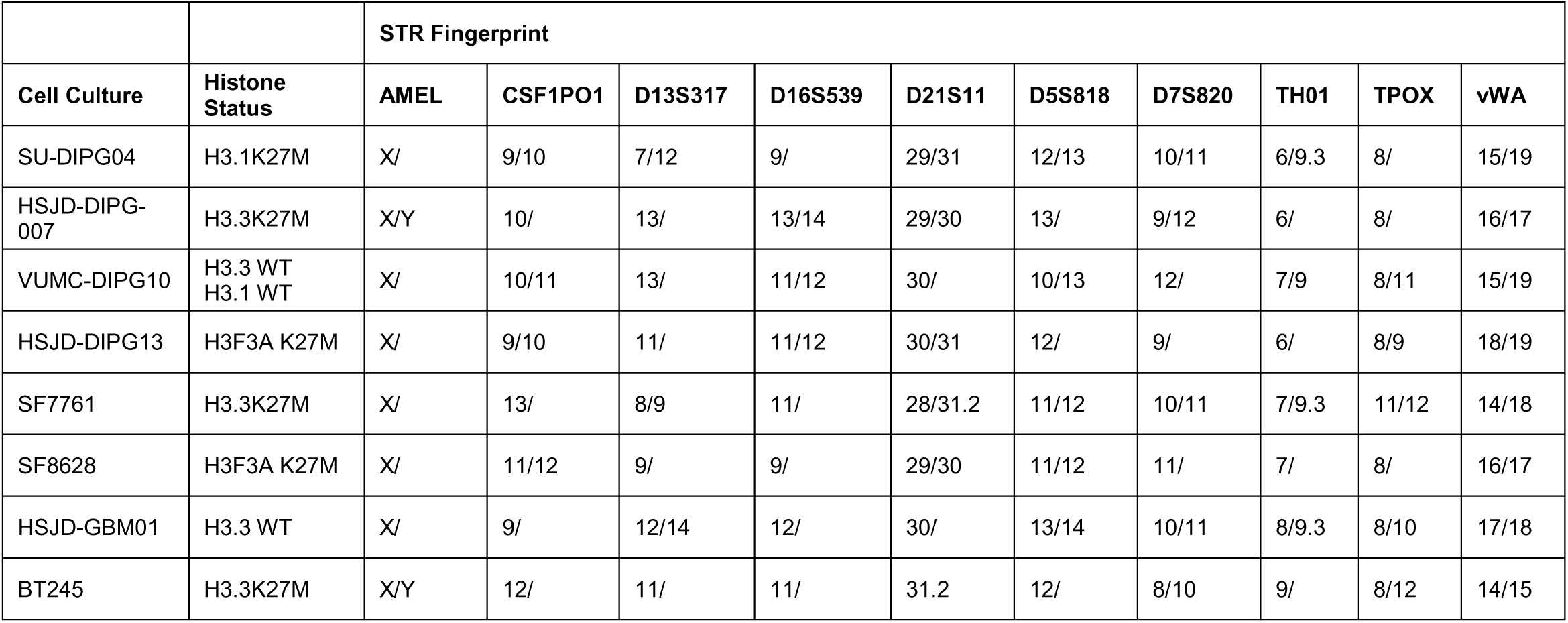
STR Fingerprinting of DMG cell lines. Cell lines validated by DNA fingerprinting through the University of Colorado Molecular Biology Service Center utilizing the STR DNA Profiling PowerPlex-16 HS Kit.

## References

1. L. M. Hoffman et al., Clinical, Radiologic, Pathologic, and Molecular Characteristics of Long-Term Survivors of Diffuse Intrinsic Pontine Glioma (DIPG): A Collaborative Report From the International and European Society for Pediatric Oncology DIPG Registries. J Clin Oncol 36, 1963–1972 (2018).

2. M. H. Jansen, D. G. van Vuurden, W. P. Vandertop, G. J. Kaspers, Diffuse intrinsic pontine gliomas: a systematic update on clinical trials and biology. Cancer treatment reviews 38, 27–35 (2012).

3. U. Bartels et al., Proceedings of the diffuse intrinsic pontine glioma (DIPG) Toronto Think Tank: advancing basic and translational research and cooperation in DIPG. Journal of neuro-oncology 105, 119–125 (2011).

4. D. N. Louis et al., The 2016 World Health Organization Classification of Tumors of the Central Nervous System: a summary. Acta neuropathologica 131, 803–820 (2016).

5. S. Nakata et al., Histone H3 K27M mutations in adult cerebellar high-grade gliomas. Brain tumor pathology 34, 113–119 (2017).

6. S. Ryall et al., Targeted detection of genetic alterations reveal the prognostic impact of H3K27M and MAPK pathway aberrations in paediatric thalamic glioma. Acta neuropathologica communications 4, 93 (2016).

7. C. Jones, S. J. Baker, Unique genetic and epigenetic mechanisms driving paediatric diffuse high-grade glioma. Nature reviews. Cancer 14, (2014).

8. D. A. Khuong-Quang et al., K27M mutation in histone H3.3 defines clinically and biologically distinct subgroups of pediatric diffuse intrinsic pontine gliomas. Acta neuropathologica 124, 439–447 (2012).

9. G. Wu et al., Somatic histone H3 alterations in pediatric diffuse intrinsic pontine gliomas and non-brainstem glioblastomas. Nature genetics 44, 251–253 (2012).

10. G. Wu et al., The genomic landscape of diffuse intrinsic pontine glioma and pediatric non-brainstem high-grade glioma. Nat Genet 46, 444–450 (2014).

11. S. Bender et al., Reduced H3K27me3 and DNA hypomethylation are major drivers of gene expression in K27M mutant pediatric high-grade gliomas. Cancer cell 24, 660–672 (2013).

12. J. Loven et al., Selective inhibition of tumor oncogenes by disruption of super-enhancers. Cell 153, 320–334 (2013).

13. J. Hagenbuchner, M. J. Ausserlechner, Targeting transcription factors by small compounds--Current strategies and future implications. Biochem Pharmacol 107, 1–13 (2016).

14. J. Shahbazi et al., The Bromodomain Inhibitor JQ1 and the Histone Deacetylase Inhibitor Panobinostat Synergistically Reduce N-Myc Expression and Induce Anticancer Effects. Clin Cancer Res 22, 2534–2544 (2016).

15. T. Sharifnia et al., Small-molecule targeting of brachyury transcription factor addiction in chordoma. Nat Med 25, 292–300 (2019).

16. H. L. Franco, W. L. Kraus, No driver behind the wheel? Targeting transcription in cancer. Cell 163, 28–30 (2015).

17. P. Filippakopoulos, S. Knapp, Targeting bromodomains: epigenetic readers of lysine acetylation. Nat Rev Drug Discov 13, 337–356 (2014).

18. I. C. Taylor et al., Disrupting NOTCH Slows Diffuse Intrinsic Pontine Glioma Growth, Enhances Radiation Sensitivity, and Shows Combinatorial Efficacy With Bromodomain Inhibition. J Neuropathol Exp Neurol 74, 778–790 (2015).

19. A. Piunti et al., Therapeutic targeting of polycomb and BET bromodomain proteins in diffuse intrinsic pontine gliomas. Nat Med 23, 493–500 (2017).

20. S. Nagaraja et al., Transcriptional Dependencies in Diffuse Intrinsic Pontine Glioma. Cancer Cell 31, 635–652.e636 (2017).

21. J. D. Larson et al., Histone H3.3 K27M Accelerates Spontaneous Brainstem Glioma and Drives Restricted Changes in Bivalent Gene Expression. Cancer Cell 35, 140–155.e147 (2019).

22. Z. Luo, C. Lin, A. Shilatifard, The super elongation complex (SEC) family in transcriptional control. Nat Rev Mol Cell Biol 13, 543–547 (2012).

23. C. Lin et al., AFF4, a component of the ELL/P-TEFb elongation complex and a shared subunit of MLL chimeras, can link transcription elongation to leukemia. Mol Cell 37, 429–437 (2010).

24. F. X. Chen, E. R. Smith, A. Shilatifard, Born to run: control of transcription elongation by RNA polymerase II. Nature reviews. Molecular cell biology 19, 464–478 (2018).

25. I. Jonkers, J. T. Lis, Getting up to speed with transcription elongation by RNA polymerase II. Nature reviews. Molecular cell biology 16, 167–177 (2015).

26. Q. Zhou, T. Li, D. H. Price, RNA polymerase II elongation control. Annual review of biochemistry 81, 119–143 (2012).

27. B. M. Peterlin, D. H. Price, Controlling the elongation phase of transcription with P-TEFb. Molecular cell 23, 297–305 (2006).

28. C. Lin et al., Dynamic transcriptional events in embryonic stem cells mediated by the super elongation complex (SEC). Genes Dev 25, 1486–1498 (2011).

29. E. Smith, C. Lin, A. Shilatifard, The super elongation complex (SEC) and MLL in development and disease. Genes Dev 25, 661–672 (2011).

30. J. E. Bradner, D. Hnisz, R. A. Young, Transcriptional Addiction in Cancer. Cell 168, 629–643 (2017).

31. K. Liang et al., Targeting Processive Transcription Elongation via SEC Disruption for MYC-Induced Cancer Therapy. Cell 175, 766–779.e717 (2018).

32. T. Fowler et al., Regulation of MYC expression and differential JQ1 sensitivity in cancer cells. PloS one 9, e87003 (2014).

33. A. M. Saratsis et al., Comparative multidimensional molecular analyses of pediatric diffuse intrinsic pontine glioma reveals distinct molecular subtypes. Acta Neuropathol 127, 881–895 (2014).

34. M. Levine, Paused RNA polymerase II as a developmental checkpoint. Cell 145, 502–511 (2011).

35. J. Zeitlinger et al., RNA polymerase stalling at developmental control genes in the Drosophila melanogaster embryo. Nat Genet 39, 1512–1516 (2007).

36. V. S. Chopra, J. W. Hong, M. Levine, Regulation of Hox gene activity by transcriptional elongation in Drosophila. Curr Biol 19, 688–693 (2009).

37. M. G. Filbin et al., Developmental and oncogenic programs in H3K27M gliomas dissected by single-cell RNA-seq. Science (New York, N.Y.) 360, 331–335 (2018).

38. Y. Chen, P. Cramer, Structure of the super-elongation complex subunit AFF4 C-terminal homology domain reveals requirements for AFF homo- and heterodimerization. J Biol Chem 294, 10663–10673 (2019).

39. F. Morales, A. Giordano, Overview of CDK9 as a target in cancer research. *Cell cycle (Georgetown*, Tex.) 15, 519–527 (2016).

40. T. Narita et al., Cyclin-dependent kinase 9 is a novel specific molecular target in adult T-cell leukemia/lymphoma. Blood 130, 1114–1124 (2017).

41. C. H. Huang et al., CDK9-mediated transcription elongation is required for MYC addiction in hepatocellular carcinoma. Genes & development 28, 1800–1814 (2014).

42. S. A. Choi et al., Identification of brain tumour initiating cells using the stem cell marker aldehyde dehydrogenase. Eur J Cancer 50, 137–149 (2014).

43. P. W. Lewis et al., Inhibition of PRC2 activity by a gain-of-function H3 mutation found in pediatric glioblastoma. Science 340, 857–861 (2013).

44. B. Krug et al., Pervasive H3K27 Acetylation Leads to ERV Expression and a Therapeutic Vulnerability in H3K27M Gliomas. Cancer Cell 35, 782–797.e788 (2019).

45. S. Nagaraja et al., Histone Variant and Cell Context Determine H3K27M Reprogramming of the Enhancer Landscape and Oncogenic State. Mol Cell, (2019).

46. An integrated encyclopedia of DNA elements in the human genome. Nature 489, 57–74 (2012).

47. J. N. Anastas et al., Re-programing Chromatin with a Bifunctional LSD1/HDAC Inhibitor Induces Therapeutic Differentiation in DIPG. Cancer Cell 36, 528–544.e510 (2019).

48. M. Malumbres, Cyclin-dependent kinases. Genome biology 15, 122 (2014).

49. G. W. Muse et al., RNA polymerase is poised for activation across the genome. Nature genetics 39, 1507–1511 (2007).

50. R. Hashizume et al., Pharmacologic inhibition of histone demethylation as a therapy for pediatric brainstem glioma. Nat Med 20, 1394–1396 (2014).

51. C. S. Grasso et al., Functionally defined therapeutic targets in diffuse intrinsic pontine glioma. Nat Med 21, 555–559 (2015).

52. G. L. Lin et al., Therapeutic strategies for diffuse midline glioma from high-throughput combination drug screening. Sci Transl Med 11, (2019).

53. A. N. Boettiger, M. Levine, Synchronous and stochastic patterns of gene activation in the Drosophila embryo. Science 325, 471–473 (2009).

54. L. J. Core, J. J. Waterfall, J. T. Lis, Nascent RNA sequencing reveals widespread pausing and divergent initiation at human promoters. Science 322, 1845–1848 (2008).

55. J. E. Karp et al., Randomized phase II study of two schedules of flavopiridol given as timed sequential therapy with cytosine arabinoside and mitoxantrone for adults with newly diagnosed, poor-risk acute myelogenous leukemia. Haematologica 97, 1736–1742 (2012).

56. M. C. Lanasa et al., Final results of EFC6663: a multicenter, international, phase 2 study of alvocidib for patients with fludarabine-refractory chronic lymphocytic leukemia. Leukemia research 39, 495–500 (2015).

57. M. A. Dickson et al., A phase I pharmacokinetic study of pulse-dose vorinostat with flavopiridol in solid tumors. Investigational new drugs 29, 1004–1012 (2011).

58. C. Fabre et al., Clinical study of the novel cyclin-dependent kinase inhibitor dinaciclib in combination with rituximab in relapsed/refractory chronic lymphocytic leukemia patients. Cancer chemotherapy and pharmacology 74, 1057–1064 (2014).

59. E. I. Heath, K. Bible, R. E. Martell, D. C. Adelman, P. M. Lorusso, A phase 1 study of SNS-032 (formerly BMS-387032), a potent inhibitor of cyclin-dependent kinases 2, 7 and 9 administered as a single oral dose and weekly infusion in patients with metastatic refractory solid tumors. Investigational new drugs 26, 59–65 (2008).

60. V. Amani et al., Polo-like Kinase 1 as a potential therapeutic target in Diffuse Intrinsic Pontine Glioma. BMC cancer 16, 647 (2016).

61. Y. Hu, G. K. Smyth, ELDA: extreme limiting dilution analysis for comparing depleted and enriched populations in stem cell and other assays. Journal of immunological methods 347, 70–78 (2009).

62. Y. Zhang et al., Model-based analysis of ChIP-Seq (MACS). Genome Biol 9, R137 (2008).

63. G. Yu, L. G. Wang, Q. Y. He, ChIPseeker: an R/Bioconductor package for ChIP peak annotation, comparison and visualization. Bioinformatics 31, 2382–2383 (2015).

